# Modelling and assessing additional transmission routes for porcine reproductive and respiratory syndrome virus: vehicle movements and feed ingredients

**DOI:** 10.1101/2021.07.26.453902

**Authors:** Jason A. Galvis, Cesar A. Corzo, Gustavo Machado

## Abstract

Accounting for multiple modes of livestock disease dissemination in epidemiological models remains a challenge. We developed and calibrated a mathematical model for transmission of porcine reproductive and respiratory syndrome virus (PRRSV), tailored to fit nine modes of between-farm transmission pathways including: farm-to-farm proximity (local transmission), contact network of batches of pigs transferred between farms (pig movements), re-break probabilities for farms with previous PRRSV outbreaks, with the addition of four different contact networks of transportation vehicles (vehicles to transport pigs to farms, pigs to markets, feed and crew) and the amount of animal by-products within feed ingredients (e.g. animal fat or meat and bone meal). The model was calibrated on weekly PRRSV outbreaks data. We assessed the role of each transmission pathway considering the dynamics of specific types of production (i.e., sow, nursery). Although our results estimated that the networks formed by transportation vehicles were more densely connected than the network of pigs transported between-farms, pig movements and farm proximity were the main PRRSV transmission routes regardless of farm types. Among the four vehicle networks, vehicles transporting pigs to farms explained a large proportion of infections, sow = 20.9%; nursery = 15%; and finisher = 20.6%. The animal by-products showed a limited association with PRRSV outbreaks through descriptive analysis, and our model results showed that the contribution of animal fat contributed only 2.5% and meat and bone meal only 0.03% of the infected sow farms. Our work demonstrated the contribution of multiple routes of PRRSV dissemination, which has not been deeply explored before. It also provides strong evidence to support the need for cautious, measured PRRSV control strategies for transportation vehicles and further research for feed by-products modeling. Finally, this study provides valuable information and opportunities for the swine industry to focus effort on the most relevant modes of PRRSV between-farm transmission.

## 1. Introduction

Porcine reproductive and respiratory syndrome virus (PRRSV) remains a major economic burden in North America (Holtkamp et al., 2013) as it continues to spread across multiple pig-producing companies (Sanhueza et al., 2019; Jara et al., 2020; Galvis et al., 2021). A recent study developed a mathematical model to reconstruct the between-farm PRRSV dynamics to reveal the role of between-farm pig movements, farm-to-farm proximity, and the continued circulation of PRRSV within infected sites (named as re-break) on PRRSV transmission (Galvis et al., 2021). Despite the promising results, the study did not fully consider indirect contacts through between-farm transportation vehicles (e.g. vehicles transporting pigs, feed or farm personnel) contact networks, which has been previously described as one of the major modes of between-farm transmission of diseases in swine (Büttner and Krieter, 2020; Porphyre et al., 2020; Niederwerder, 2021), such as PRRS (Dee et al., 2002, 2004; Pitkin et al., 2009; Thakur, Sanchez et al., 2015) and African swine fever (ASF) (Gao et al., 2021; Gebhardt et al., 2021).

Detailed data regarding transportation vehicle movement and routes, coming in and out pig premises, can be difficult to obtain, which indeed could help explaining the lack of models considering this transmission pathway (Bernini et al., 2019; EFSA Panel on Animal Health and Welfare (AHAW) et al., 2021). Previous studies which have approached indirect transmission by transportation vehicles, have either used simulated probabilities to define indirect contact between farms such as the studies by Thakur et al., 2015 and Wiltshire, 2018, or observed truck movements such as the study by Buttner, 2020 using between farms movements in Germany. The potential of viral stability on vehicle surfaces has been demonstrated to be dependent on environmental conditions such as temperature, pH, moisture, and vehicle disinfection procedures (Dee et al., 2002, 2003; Jacobs et al., 2010). For example, Dee et al., isolated PRRSV under field conditions from surfaces such as concrete, floor mats and fomites between 2h and 4h after surface contamination (Dee et al., 2002). PRRSV has been shown to be more stable at low temperatures (−20° C to −70° C), surviving for long periods of time (>4 months) (Benfield et al., 1992), and becoming unstable as the temperature increases (Jacobs et al., 2010). In addition, dry conditions, low pH ranging between 5 and 7 (Benfield et al., 1992), iodine, quaternary ammonium or chlorine compounds used in vehicle disinfection were successful in inactivating PRRSV (Shirai et al., 2000). Thus, if environmental conditions favor PRRSV survivability and the vehicles are poorly cleaned and disinfected, the potential for pathogens to disseminate within highly connected networks could indeed play a major role in disease spread (Büttner and Krieter, 2020; Gebhardt et al., 2021).

In addition to transportation movements, contaminated feed could represent a possible route for pathogen transmission (Gebhardt et al., 2021; Niederwerder, 2021), but the probability of PRRSV transmission through feed has been described as relatively low (Cochrane et al., 2017; Ochoa et al., 2018; Blázquez et al., 2020). However, a recent report by Dee et al. 2020, demonstrated under experimental conditions that pigs which consumed pellet feed contaminated with 1×10^5^ TCID50/ml of PRRSV became infected. As such, cross-contamination of pellet feed may occur when coming into direct contact with contaminated fomites or feed mill workers after the pelleting process (Niederwerder, 2021). In addition, it is important to acknowledge that inadequate temperature applied during the pelleting process could reduce the probability to inactivate PRRSV from contaminated feed ingredients, as observed in an experimental study with porcine epidemic diarrhea virus (Cochrane et al., 2017). In North America, most feed formulations include some animal by-products, such as animal fat, dried plasma, or bone meal in order to increase growth performance (Lewis and Southern, 2001). Without adequate processes to inactivate PRRSV, these ingredients could potentially be a source of contamination and later infection. Magar and Larochellle in 2004 surveyed two Canadian slaughterhouses and found 4.3% of animal serum samples and 1.2% of the meat samples were positive for PRRSV by polymerase chain reaction. The same study also demonstrated that the consumption of contaminated animal by-products caused infection in pigs (Magar and Larochelle, 2004).

In this study, we built a novel mathematical model of PRRSV transmission tailored to nine modes of between-farm propagation: local transmission by the farm-to-farm proximity, between-farm animal and re-break for farms with previous PRRSV outbreaks (Galvis et al., 2021), with the addition of between-farm vehicle movements (feed, shipment of live pigs between farms and to slaughterhouses, and farm personnel [crews]), and the quantity of animal by-products which was restricted to animal fat, meat and bone meal in pig feed ingredients. The model was used to estimate the weekly number of new PRRSV outbreaks and their spatial distribution, which were compared to available data, and to quantify the contribution of each transmission route.

## Material and Methods

### Databases

In this study, we used weekly PRRSV records captured by the Morrison Swine Health Monitoring Program (MSHMP) (MSHMP, 2020). Data included outbreaks between January 22, 2009 and December 31, 2020, from 2,294 farms from three non-commercially related pig production companies (coded as A, B, and C) in a U.S. region (not disclosed due to confidentiality). Additional details about the study population and PRRSV classification in the U.S. are available in (Galvis et al., 2021, Holtkamp et al., 2011). Individually, each PRRSV record was classified as a new or recurrent outbreak according to the time between consecutive outbreaks per farm (Galvis et al., 2021). A list of pig farms was available from the MSHMP database (MSHMP, 2020), which included individual national premises identification number, farm type (sow [which included farrow, farrow-to-wean, and farrow-to-feeder farms], nursery, finisher [which included wean-to-feeder, wean-to-finish, feeder-to-finish], gilt development unit [which could be either part in finisher or sow farms depending upon farm type used by pig production company], isolation and boar stud), pig spaces per farm, and geographic coordinates. Between-farm pig movement data from January 01, 2020, to December 31, 2020, were used to reconstruct directed weekly contact networks. Each movement batch included movement date, farm of origin and destination, number of pigs transported, and purpose of movement (e.g., weaning). Movement data missing either the number of animals transported, farm type, farm of origin, or destination were excluded prior to analysis (731 [1.2%] unique movements were excluded). In addition, four networks formed by transportation vehicles were recorded from the global positioning system (GPS) vehicle tracker for which included tracking of all farms from company A (76% of all farms within the study region for 2020) from January 01, 2020, to December 31, 2020. These movements comprise near-real time GPS records of each transporting vehicle, which include geographic coordinates for every 5 seconds, of any vehicle. Overall, 398 vehicles were monitored which included: (i) 159 trucks used to deliver feed to farms, (ii) 118 trucks utilized in the transportation of live pigs between farms, (iii) 89 truck used in the transportation of pigs to markets (slaughterhouse), and (iv) 32 vehicles used in the transportation of crew members, which by the information we collected correspond to the movement of additional personnel needed for vaccination, pig loading and unloading among other activities which included power washing (Figure 1). Each movement batch included a unique identification number, speed, date and time along with coordinates of each vehicle location recorded every 5 seconds. A vehicle visit was defined as a vehicle coordinate (latitude and longitude) and speed of zero km/h for at least 5 minutes within a radius of 1.5 km of any farm or cleaning station (time and distance radius selected after discussion with personnel in charge of vehicle logistics and data observation). In case more than one farm was located within a 1.5 km radius, we assumed all these farms were at-risk of transmission, and therefore the vehicle contacted all of them. It is worth noting that in some cases where farms are located at distances <1.5 km from other farms it is because they belong to the same pig production (personal communication). We calculated the time in minutes the vehicle remained within each farm’s perimeter and the vehicle contact networks between the farms was built considering the elapsed time a vehicle visited two or more different farms (Figure 1). To accommodate PRRSV survivability in the environment, we considered two seasons (cold and warm weather) based on previous literature (Dee et al., 2002; Jacobs et al., 2010). Under laboratory conditions, it was reported that PRRSV preserved stability for more than 72h when temperatures oscillated between 4° C and 10° C (cold temperatures) and less than 24h when temperatures were equal or higher than 20° C (warm temperatures) (Jacobs et al., 2010). Thus, an edge among two different farms was recorded if the elapsed time the vehicle visited both farms was less than 72 hours or 24 hours, for the cold and warm seasons, respectively. However, we did not consider the formation of edges between consecutive farms after vehicles were observed via GPS driving through clean stations (Figure 1). The edges for all four vehicle networks were weighted by the elapsed time each vehicle visited two different farms, which was later transformed to a probability assuming a decreasing linear relationship of PRRSV stability in the environment (Figure 1 and Supplementary Material Figure S1). Additionally, we collected feed load out records from all (three) feed mills of company A for 2020, with each feed record including feed mill identification with individualized feed formulation (ingredients), amount of feed delivered, destination farm identification, and destination farm delivery data. From the feed records, we collected the amount in pounds (lb) of animal by-products (parts of a slaughtered animal that included animal fat, pig plasma and meat and bone meal) of each feed formulation received by the farms for each week of 2020 (Supplementary Material Figure S2 and S3). Although company B and C data about vehicle movements and feed delivery was not available, we kept the farms from both companies in the transmission model to complement the PRRSV dissemination by the local transmission (Jara et al., 2020).

**Figure 1.**
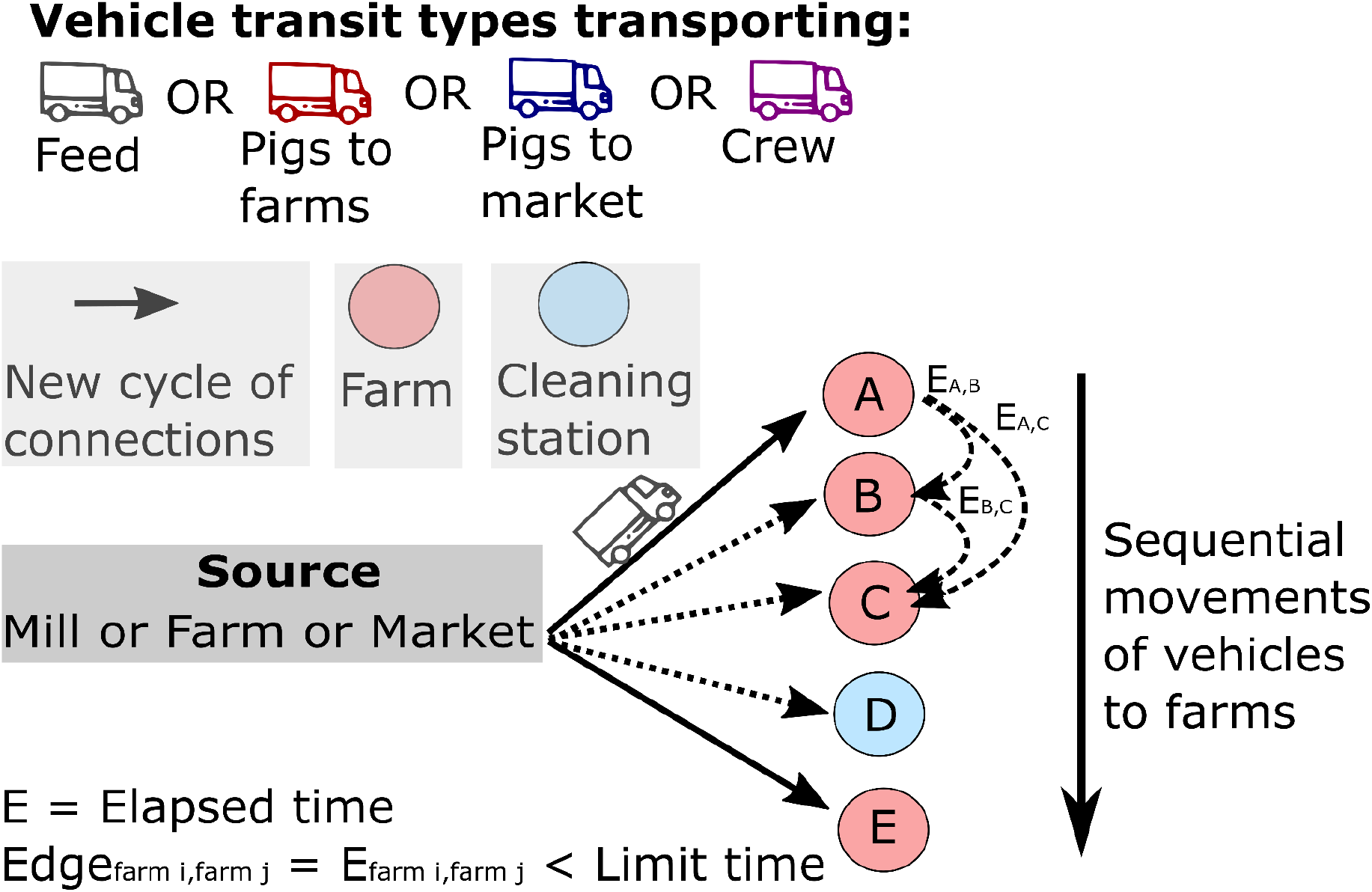
Framework of the indirect farm contacts formed by transportation vehicle movements. The transportation vehicle networks were reconstructed based on consecutive farm visits of each vehicle. Because the stability of PRRSV in the environment is directly impacted by the environmental conditions, a contact network was reconstructed considering all edges between farm visits that happened within 72 hours for cold months (from October until March) and within 24 hours for warm months (from April until September).

### Descriptive analysis

#### Between farm animal and transportation vehicles movement

The networks formed by movement of live pigs transported between farms and four types of transportation vehicles visiting farms were reconstructed and analyzed. A set of network metrics including: size, general properties, and heterogeneity at node-level were evaluated for each directed static and temporal network (Supplementary Material Table S1 for terminology and network metric description). To determine if the static networks of pig and vehicle movements could represent the temporal variation of the between farm contacts over a year, we calculated the causal fidelity (Lentz et al., 2016). Briefly, causal fidelity quantifies the error of the static representation of temporal networks through a ratio among the number of paths between both representations. Thus, a casual fidelity of 100% would mean that a temporal network is well represented by its static counterpart, and conversely, a value close to 0% means the network should not be considered as a static system (Lentz et al., 2016). We then estimated whether farms with PRRSV outbreaks records were more frequently connected with other infected farms through the ingoing and outgoing contact chain, compared with farms without PRRSV records. Finally, we estimated if the time transportation vehicles remained within farms premises was higher in the farms with PRRSV outbreaks compared with farms without PRRSV breaks. The association for the contact chain and the time the vehicles stayed in the farms with PRRSV outbreaks were evaluated through a Mann-Whitney test.

#### Animal by-products in feed ingredients

We calculated the total amounts of animal fat, pig plasma, animal protein blend 58%, protein blend (animal protein blend and protein blend was made of a combination of ingredients such as meat meal, corn germ meal, hominy and dried distillers grains with solubles), and meat and bone meal present in each of the 23 feed formulations delivered to farms with and without PRRSV outbreaks in 2020 (Supplementary Material Figure S2 and S3). In order to further evaluate the association between PRRSV outbreaks and the delivery of feed with animal by-products, we performed a logistic regression analysis for each farm type and ingredient in which the response variable was positive or negative for PRRSV from January 1^st^, 2020 to December 31^st^, 2020, and the predictor was the amount of animal by-product divided by the farm’s pig capacity to avoid confusion by the farm size.

### Transmission model

The analysis of spatiotemporal distribution of farm-level PRRSV outbreaks was based on our previously developed stochastic model (Galvis et al., 2021), which here was extended to include vehicle transportation networks and the delivery of animal by-products. The model was calibrated on the weekly PRRSV outbreaks and considering nine transmission modes including: (1) contact network of discrete pig movements; (2) the local transmission events between neighboring farms driven by distances among farms; (3) re-break by a previous exposure to PRRSV; indirect contact by vehicles coming into farms, including for (4) feed, animal delivery to (5) farms and (6) market, and (7) vehicles used by personnel (crew) involved in the loading and unloading of pigs; amount of (8) animal fat and (9) meat and bone meal in feed formulation delivered to farms (Figure 2). The model simulates between farm transmission among three farm-level infectious states, Susceptible-Infected-Outbreak (SIO), and we defined Susceptible status as farms free of PRRSV, Infected status as farms with PRRSV but not yet detected, and Outbreaks status as infected farm that detected PRRSV. Thus, farms in a susceptible state (*i*) receive the force of infection of infected and outbreak farms (*j*) in each time step *t* and become infected at probability *Y_it_* (Figure 2). It is worth noting that the latent period of PRRSV is not explicitly modeled, as it is typically a few days after infection, and often viral shedding starts within 7 days post-infection (Pileri and Mateu, 2016; Chase-Topping et al., 2020), thus it is embedded in the weekly timestep. Local transmission was modelled through a gravity model where the probability of infection is proportional to the animal capacity of the farms and inversely related to the distance between the two farms (i.e. lower transmission at longer distances), with the maximum distance set at 35 km, similar to our previous study to facilitate the comparison of results (Galvis et al., 2021). Local transmission is also dependent on the enhanced vegetation index (EVI) around the farm *i* (Jara et al., 2020; Galvis et al., 2021), such that the probability of transmission decreases with high EVI values (Supplementary Material Figure S4). The transmission associated with between farm pig movements is modeled by the number of all infected and outbreak farms sending pigs to susceptible farms. The dissemination via transportation vehicle networks (e.g. vehicles transporting pigs to farms) is modeled by the edge weight (E) and the time the vehicles remained on the susceptible farm premise (*Z_it_*) (Figure 1 and Supplementary Material Figure S1 and S5). The transmission via animal fat and meat and bone meal was only considered to sow farms and modulated simply by the amounts delivered to susceptible farms (*A_it_*). Pig plasma, animal protein blend meal 58% and protein blend were only delivered to nursery and finisher farms, thus were not considered. For the re-break rate which is only considered for sow farms, we assumed that subsequent new infections at an individual farm, within a time period of two years, were associated with the same strain as the previous outbreak, and the probability was based on a survival analysis evaluating the time farms re-break after recovering (*W_it_*) (Holtkamp et al., 2010) (Supplementary Material Figure S6). Then, for each transmission route, the force of infection (*λ*) of infected and outbreak farms varies with a seasonality derived from analysis of the PRRSV records from 2015 to 2019 (Supplementary Material Figure S7). In addition, sow farms without a record of PRRSV outbreaks since 2009 were assumed to have high biosecurity levels (*H*) that reduce the force of infection received by infected and outbreak farms, being *H* higher than zero and calibrated according to the observed outbreaks (Supplementary Material Table S2). Otherwise, farms with outbreaks records were assumed to have low biosecurity levels and *H* was defined as zero.

**Figure 2.**
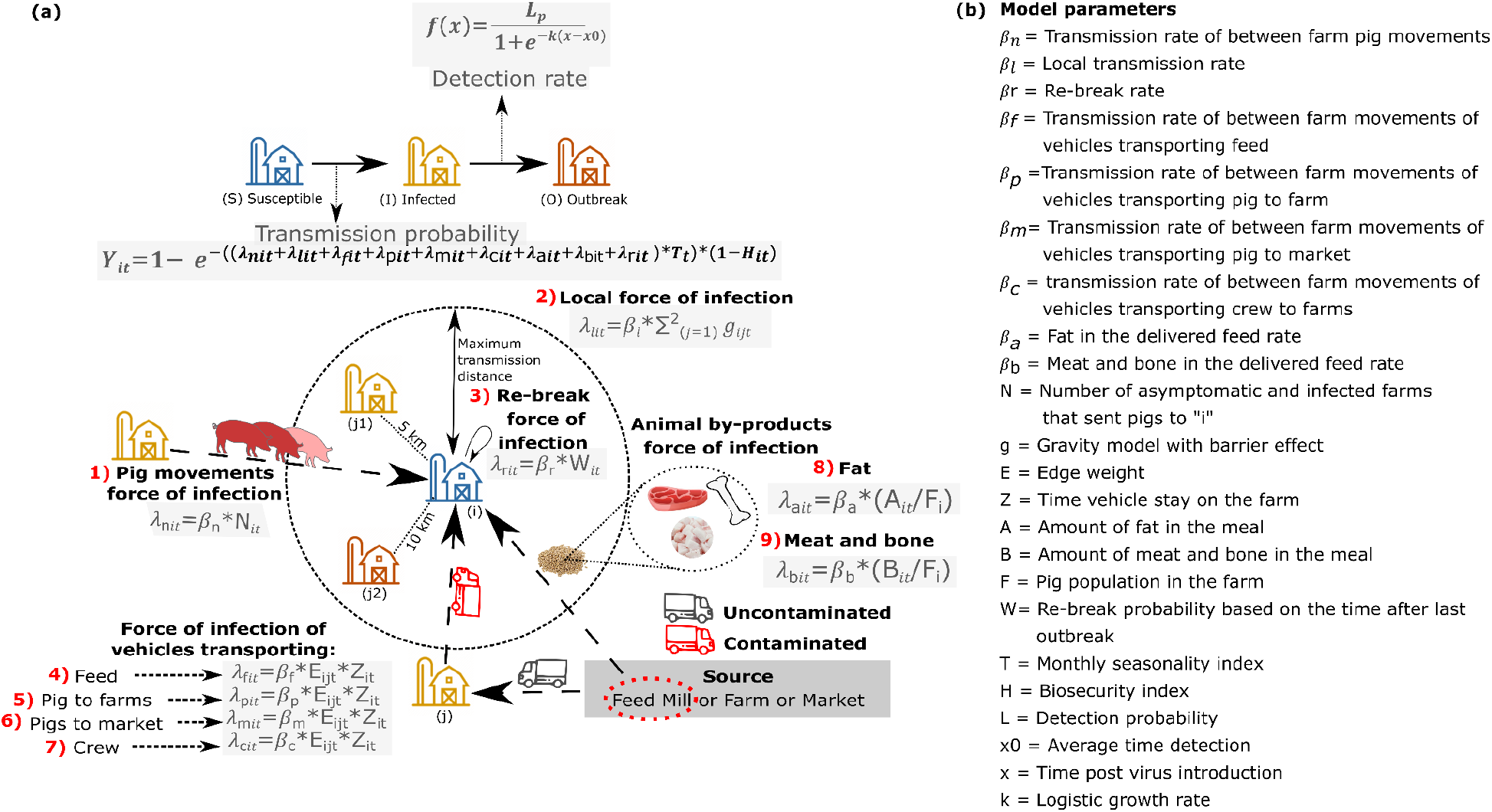
Transmission model framework. (a) Model flowchart of the farm’s infectious status and routes of PRSV transmission, and (b) description of the model transmission parameters. In the example, for the local infection force, j takes the values from 1 and 2 because these are the only two farms within the maximum radius distance.

The transition from infected to an outbreak farm is estimated through a detection rate *f(x)* (Figure 2). Thus, the probability that farms transit to outbreak state is assumed to be dependent on the maximum detection probability (*L*), considered equal to cases reported to MSHMP (MSHMP, 2020), and the average time it takes a farm to detect the disease (*x0*), assumed to be 4 weeks (estimated from information provided by local swine veterinarians and previous literature) (Neira et al., 2017). The proportion of Infected and Outbreak sow farms that return to a susceptible state is drawn from a Poisson distribution with mean of 41 weeks, which is the average time to stability described elsewhere (Sanhueza et al., 2019). Nursery and finisher farms’ transition to susceptible status are driven by pig production movement scheduling of the all-in all-out management schemes of closeouts or by incoming or outgoing movements, whichever came first (Galvis et al., 2021). Briefly, nurseries and finisher farms become susceptible within 7 and 25 weeks of pig placement, respectively, or when at least one new pig movement is recorded before the farm reaches the scheduled production phase timeline described earlier. A detailed description of the model can be found in Figure 2, and previous work describes in greater detail other model parameters (Galvis et al., 2021). Finally, we used an Approximate Bayesian Computation (ABC) rejection algorithm (Minter and Retkute, 2019), to estimate the posterior distribution of unknown model parameters (list of model parameters available in Supplementary Material Table S3) by selecting the 100 particles (number of particles accepted defined according to our computational resources) that best fitted the temporal and spatial distribution of observed PRRSV outbreaks (Supplementary Material Figures S8-S10).

### Model outputs

The model outputs included (a) the force of infection for each farm type and transmission route, (b) the weekly number of infected undetected and detected farms (outbreaks) and (c) the sensitivity performance to detect PRRSV outbreak locations (Supplementary Material Section 2). We carried out 1,000 simulations, each simulation choosing at random one of the 100 accepted particles, to estimate the relative contribution of each transmission route and the weekly number of cases. For the sensitivity performance, 100 simulations were used for each particle to estimate the risk of infection, and only 100 particles were used due to computational resources (additional information see Supplementary Information Figure S10). For the contribution of the routes, we evaluated the number of infected farms resulting from each transmission route individually, which were then divided by the number of simulated infected farms from all the combined routes and commercial companies. In addition, the average contribution for each route was estimated by summing the weekly contributions divided by the number of simulated weeks, and credible intervals (CI) using an equal-tailed interval method were estimated from the weekly contribution distribution. The model was developed in the R (3.6.0 R Core Team, Vienna, Austria) environment, and all simulations were run in RStudio Pro (1.2.5033, RStudio Team, Boston, MA) and transmission model framework is available at https://github.com/machado-lab/pigspread.

## Results

The comparison among the five networks showed that vehicles transporting feed, pigs to farms, pigs to markets and movement of crews were significantly more connected than the pig movement network (Table 1). The network density formed by vehicles transporting feed exhibited the highest density (edge density = 0.2). Comparing the paths between the static and temporal networks of pig movement and transportation vehicles movements, we found that causal fidelity was above 32% for all networks (Table 1), which means that a significant number of the causal paths described by the static networks can be found in the temporal networks. The networks of vehicles transporting feed and pigs to farms exhibited the highest causal fidelity values (causal fidelity >90%), which means these networks are the closest representation of causal paths between the static and temporal network. Analyzing the network components of each vehicle network for company A, we found that the Largest Strongly Connected Component (LSCC) had between 976 and 1,591 farms, thus these vehicle movement networks connected between 55% and 90% of company A farms (Table 1). For the pig movement network, company B showed the highest number of farms in the LSCC with 61 farms, which represented 27% of the farms from that company, while company A and C showed a low level of connectivity with the LSCC representing less than 1% of the farms. Comparing pig movement networks also showed that company B had the highest edge density of 0.013, followed by company C (edge density of = 0.005) and company A the lowest (edge density of = 0.002).

**Table 1.**
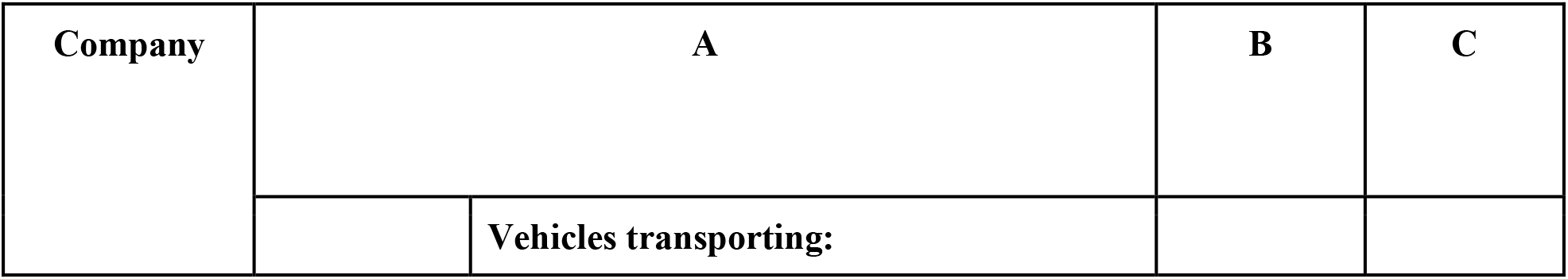
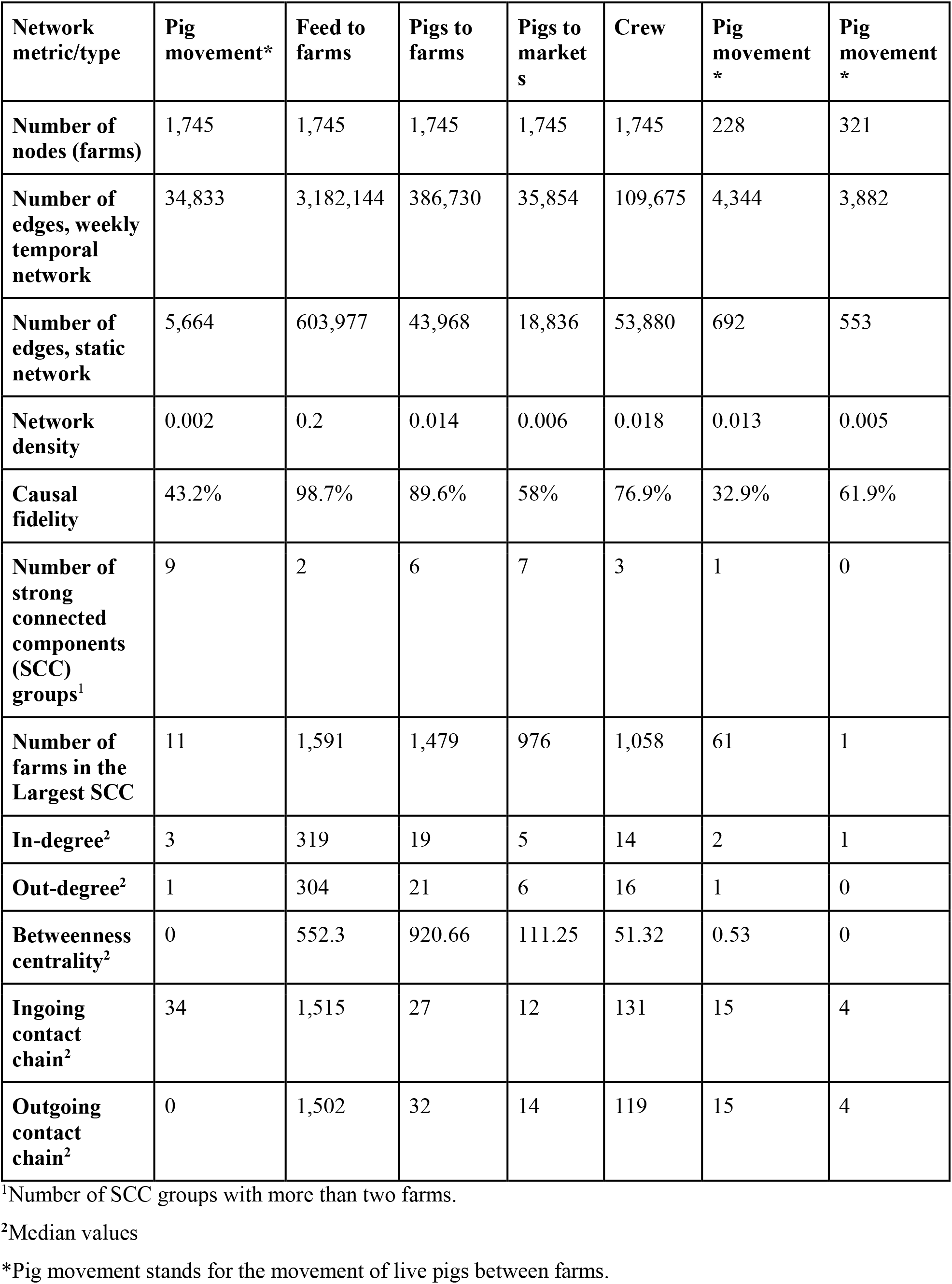
Summary of the network metric of pig movements and vehicle movements of the three different pig producing companies, with data from January 2020 until December 2020.

The vehicles transporting feed static network had a median in-degree and out-degree of 319 and 304, respectively, which was the highest in comparison with vehicles transporting pigs to farms, pigs to markets and vehicles used in the transportation of crew (Table 1, Supplementary Material Figure S11 and S12). The networks of vehicles transporting pigs to farms and crew had a median in-degree and out-degree ranging between 14 and 21, while vehicles transporting pigs to markets and pig movements for all three companies showed median in-degree and out-degree less than 7 (Table 1). The network of vehicles transporting pigs to farms showed the highest median betweenness centrality, followed by vehicles transporting feed, vehicles transporting pigs to markets and then crew networks, whilst pig movements had the lowest betweenness (Supplementary Material Figure S13). Interestingly, despite the network of vehicles transporting feed being more densely connected, the network of vehicles transporting pigs to farms showed the highest median betweenness centrality values, indicating that this network has the highest number of shortest paths by farm to connect other farms (Table 1 and Supplementary Material Figure S13). Considering the result from the temporal networks, vehicles transporting feed showed the highest median ingoing (ICC) and outgoing contact chain (OCC) (Supplementary Material Table S1), which indicates that feed routes create the largest sequential paths over time that allow for more connections between farms than any transportation or pig movements network (Table 1 and Supplementary Material Figure S14 and S15). The ICC and OCC from vehicles transporting crew showed the second highest values, followed by vehicles transporting pigs to farms and then vehicles transporting pigs to markets. When considering pig movement, company A had the highest median ICC among the different companies (ICC = 34), and a median OCC of zero. Companies B and C showed lower ICC values, but a higher median OCC of 15 and 4, respectively (Table 1 and Supplementary Material Figure S14 and S15).

Furthermore, we evaluated the association of PRRSV outbreaks frequency within both ICC and OCC from infected and non-infected farms. The results showed that PRRSV infected farms were more frequently found within the ICC and OCC of infected farms for the networks of vehicles transporting feed, pigs to farms and pigs to markets networks (*p* < 0.05), while such association was not observed for vehicles transporting crew, and for pig movement networks it was only significant for OCC (Supplementary Material Figures S16 and S17). We also evaluated the association between the time transportation vehicles remained within infected and non-infected farms. The vehicles transporting feed and crew members spent more time within infected nursery farms when compared with the non-infected farms (*p* < 0.05), while no association was found with PRRSV at sow and finisher farms for these vehicles (*p* > 0.05) (Supplementary Material Figures S18). In addition, no differences were found among the time spent on infected farms compared with the non-infected farms for the vehicles transporting pigs to farms and pigs to markets for any farm type (*p* > 0.05). Finally, the amount of animal by-products in feed (animal fat, pig plasma, protein blend and meat and bone meal) were not significantly associated with PRRSV outbreaks (*p* > 0.05) (Supplementary Material Figures S19-S21).

From the weekly model simulations from January 1th until December 31th, 2020, we estimated the total number of infected farms at 1,790 (95% CI: 1,776–1,804), 113 (95% CI: 112–113) of which corresponded to infected sow farms, 715 (95% CI: 704–726) to nursery farms and 960 (95% CI: 952–967) to finisher farms (isolation and board stud farm were excluded from the results because no outbreaks were reported in the studied period). It is worth noticing that, just as in the data, in our simulations the same farm could have been at infected state more than once over the simulated year. Overall, results showed a good agreement between the weekly observed number of PRRSV outbreaks and simulated outbreaks (Supplementary Material Figure S9). The model inferred that, at the end of the 52 weeks, on average 158 (8.8%) of all PRRSV infected farms would be detected, in which 90% of the infected sow farms were detected, while a much lower proportion of detection was estimated for nurseries (4.8%) and finishers farms (2%). The model’s predictive performance to correctly identify the weekly spatial distribution of known PRRSV outbreaks showed an area under the ROC curve of 0.7 (More details are available in Supplementary Material Section 2).

Evaluating the contribution of nine transmission routes over the simulated PRRSV spread for company A’s farms demonstrated that for sow farms the most important route was the local transmission contributing to an average of 32.4% (95% CI 15%-67%) of the farm infections, followed by pig movements with 28.3% (95% CI 1.9%-68%), vehicles transporting pig to farms with 20.9% (95% CI 5%-45%), vehicles transporting feed 12% (95% CI 0,5%-32%), re-break 3.2% (95% CI 0%-6%), amount of animal fat within feed formulation 2.5% (95% CI 0,7%-6%), vehicles transporting pigs to markets 0.4% (95% CI 0%-2.5%), vehicles transporting crew 0.2% (95% CI 0%-1.5%) and amount of meat and bone meal within feed formulation 0.03% (95% CI 0%-0,5%) (Figure 3). For nursery farms, pig movements were the most important route contributing to 76.4% (95% CI 57%-89%) of the farm infections, followed by vehicles transporting pigs to farms 15% (95% CI 5%-26%), local transmission with 5.8% (95% CI 3%-16%), vehicles transporting feed 2.3% (95% CI 0.3%-6%), vehicles transporting pigs to markets 0.44% (95% CI 0%-1.4%) and vehicles transporting crew 0.1% (95% CI 0%-0,5%). For finisher farms, local transmission was also the most important route contributing to 35.5% (95% CI 17%-58%) of the farm infections, followed by pig movements with 30.1% (95% CI 12%-52%), vehicles transporting pigs to farms 20.6% (95% CI 7%-38%), vehicles transporting feed 9.2% (95% CI 3%-19%), vehicles transporting pigs to markets 3.8% (95% CI 0.13%-11%) and vehicles transporting crew 0.61% (95% CI 0.01%-1.8%). As transportation vehicle data were not available for companies B and C, the results were restricted to three transmission pathways (pig movements, local transmission and re-break) and are available in Supplementary Material section 4, Figure S22.

**Figure 3.**
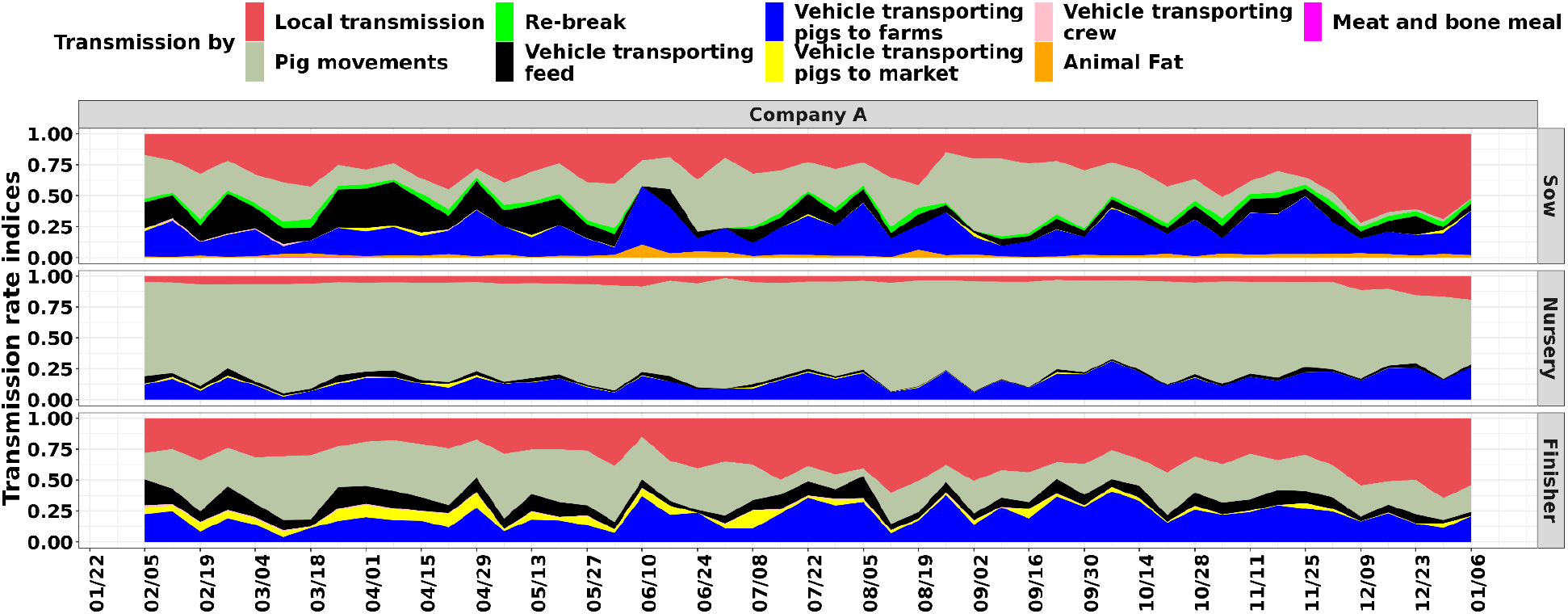
Farm infection contribution for each transmission route of each farm type (rows). The *y*-axis represents the proportion of transmission by each transmission route, while the x-axis shows each week in the simulation. Weekly proportions of transmission were calculated by dividing the number of simulated infected farms for the total number of routes combined.

## Discussion

In this study, we demonstrated the contribution of nine pathways in PRRSV dissemination dynamics which included pig movements network, farm-to-farm proximity, different types of transportation vehicle networks (vehicles transporting feed, pigs to farms, pigs to markets, and crew), the delivery of animal by-products, in particular animal fat and meat and bone meal in the feed meal and re-break. We demonstrated that the transportation vehicle networks connected more farms than the pig movement networks, therefore between farm contacts by transportation vehicles have the potential to propagate PRRSV to more sites and more quickly than moving live pigs between-farms. Thus, we remark that the greater potential for PRRSV transmission via transportation vehicle networks pose a great challenge to surveillance and effective control of endemic disease in North America and future eradication of possible introduction of foreign animal diseases such as ASF (Brown et al., 2020; Gao et al., 2021). Our regression results also suggest that the overall volume of animal by-product delivery to the farms was not associated with the PRRSV outbreaks, and our mathematical model results indicated that animal fat and meat and bone meal delivered to sow farms combined contributed to <2,6% farm infection in the simulations.

Our study addresses major gaps in the understanding of how PRRSV propagates between-farms by modelling multiple modes of transmission, which expand the understanding of PRRSV propagation, thus far restricted to studies that have only considered the dissemination of PRRSV throughout animal movement networks (Thakur, Revie et al., 2015; Lee et al., 2017; Makau et al., 2021). Our previous study (Galvis et al., 2021) and the information about the transmission dynamics of PRRSV (Dee et al., 2002; Perez et al., 2015; Thakur, Revie et al., 2015; Pileri and Mateu, 2016; Silva et al., 2019; Jara et al., 2020) highlighted limitations of model accuracy and paucity evaluation of PRRSV dissemination by not considering well known modes of transmission, such as indirect contacts formed by transportation vehicles. In this study, we have further extended the previous transmission model (Galvis et al., 2021), by including six additional transmission routes, including the contact networks formed by transportation vehicles and animal by-products delivered to sow farms via feed ingredients. Our results demonstrate that pig movements and local transmission were the main transmission routes, regardless of farm types (sow, nursery, and finisher) (Figure 3). However, the contribution of transportation vehicles used to transfer pigs to farms explained a significant number of infected farms as follows, 20.9% of sow farms, 15% of nurseries and 20.6% of finisher farms. Our examination of feed delivery to farms, more specifically the volume of animal by-products animal fat and meat and bone meal, indicated that this mode did not contribute significantly to PRRSV transmission, contributing to only 2.5% and 0.03% of the infection in sow farms, respectively. Although our results tend to prove a small role of feed, the approach used in this study is limited by several factors that affect the contribution of feed in the PRRSV dissemination, therefore these results must be taken with caution (more details in a section below). Finally, for companies B and C, the dominant routes of transmission were similar to the results for company A, with pig movements having the greatest contribution to infection at nursery farms, while local transmission mainly affected finishers (Figure 3). However, re-break contributed to a high number of infections that was not observed in our previous study (Galvis et al., 2021). Overall, our results have also reinforced the findings from previous studies which highlight the role of vehicles in the transmission of infectious diseases that pose a significant threat to or have a particular impact on the swine industry (Dee et al., 2003, 2004, 2007; Melmer et al., 2020) and the potential risk of contaminated animal by-products in the feed meals to contribute to between-farm transmission of PRRSV (Dee et al., 2020; Niederwerder, 2021).

From individual vehicle GPS movement data, we reconstructed networks while considering the elapsed time between farm visits, and the time vehicles spent within each farm to define effective contacts among them. In addition, we considered contact between farms to end once a truck drove through a cleaning station (Figure 1), as previous studies suggest that vehicle disinfection can reduce the probability of pathogens’ introduction into pig farms, e.g., PRRSV (Thakur et al., 2017) and ASFV (Yoo et al., 2021). In the current work, the inclusion of cleaning stations reduced around half the edges of the four types of transportation vehicle networks, which represent valuable information for the analysis of disease transmission given that including or not cleaning stations provide different results of the pathogen dissemination among the farms through vehicles. It is worth to notice that we assumed that the cleaning process was always effective to inactivate PRRSV (Shirai et al., 2000), even though the literature about probability of PRRSV survival on vehicle surfaces after cleaning and disinfection remains limited (Dee et al., 2004). Thus, further studies evaluating the presence and infectivity of PRRSV after vehicle cleaning and disinfection are necessary.

Previous studies attempting to model transportation vehicle networks were limited to either static or simulated between-farm contact networks, thus limiting our ability to further compare our networks results (Thakur, Sanchez et al., 2015; Wiltshire, 2018; Sterchi et al., 2019; Porphyre et al., 2020; Yang et al., 2020). However, our result demonstrated that for the four vehicle networks described in this study, the static networks provided a close representation of the actual temporal networks, in which the ratio of the number of path between the static and temporal network varied from 58% for vehicles transporting pigs to markets to 98.7% for vehicles delivering feed (Table 1). Because of the high fidelity of the vehicle networks, our results demonstrate that the static networks may be sufficient to explain causal paths among farms formed by vehicle movements (Lentz et al., 2016). Although these results were only evaluated by comparing the static and temporal network from a single commercial company and with one year of GPS data, our results strongly suggest that static views of transportation vehicles networks can be used either to describe such networks or to be used in disease transmission models when researchers only have access to a static view of a network.

The transportation vehicle networks were more densely connected than the networks of between-farm pig movements, ranging from 3 to 100 times more connected considering the network of vehicles transporting pigs to markets and feed, respectively. Other studies have also found that vehicle movements between-farms used for animal hauling increased the indirect contacts among farms by more than 50% (Porphyre et al., 2020), consequently the network weak connected component became 50% larger (Sterchi et al., 2019). Among the transportation vehicle networks analyzed here, the network of feed deliveries was the most connected; for instance more than 85% of all farms of company A were connected through sequential paths in the temporal network (ICC and OCC Table 1). On the contrary, the network of trucks transporting live pigs between-farms was less connected, but in our transmission model it represented between 15% and 20% of the total transmissions. In turn, the probability of transportation vehicles being contaminated with PRRSV is more likely in vehicles transporting animals than feed, mainly because of the direct contact with infected animals. Vehicles transporting pigs to markets could in the same way play a similar risk of contamination as vehicles transporting pigs to farms, however the former vehicles tend to visit mostly finisher farms with pigs ready for slaughter, thus less likely to re-introduce PRRSV back into the transmission chain by visiting breeding farms (Passafaro et al., 2020). It is worth noting that this is not the case for farms with complete production cycles, such as farrow to finisher farms, in which movements returning from slaughterhouses are likely to pose a great risk (Henry et al., 2018). Our study was also the first to collect and consider in a transmission model the contact networks of vehicles transporting crew between-farms. It is known that the number of farm visitors increase the probability of introduction of infectious pathogens (e.g. PRRSV), thus the movement of additional personnel often involved in loading and unloading pigs have been previously associated with PRRSV dissemination and outbreaks (Dee et al., 2002; Pitkin et al., 2009; Rossi et al., 2017). It is worth noting that in our study we modeled the consecutive contacts between farms formed by the vehicles used by crew as a route of PRRSV dissemination, by evaluating individual vehicles connecting farms that were visited consecutively rather than the group of crew members, which is known to vary between each farm visit. Therefore, the risk of infection by vehicles transporting crew could be more related to the vehicle itself as fomite, which in turn is likely to represent a lower risk of infection compared to the risk of infection related to the personnel. In general, all four transporting vehicle networks evaluated in this study contributed to PRRSV transmission, similarly to previous studies (Thakur, Sanchez et al., 2015; Rossi et al., 2017; Porphyre et al., 2020; Yang et al., 2020). It is important to remark that the contribution of the networks described above may have been underestimated, mainly because we have not fully considered the effectiveness of cleaning and disinfection to reduce PRRSV contamination in vehicles driving through cleaning stations. The lack of studies measuring the effectiveness of cleaning and disinfections stations in reducing PRRSV and the contribution of additional on-farm biosecurity cleaning and disinfection such as the presence of automatic cleaning stations at the entrance of farms (Dee et al., 2004, 2007; Silva et al., 2019), is clearly needed in order to better specify the reduction of such procedures in the force of PRRSV transmission.

The potential propagation of infectious diseases within feed ingredients has been of concern not only to the swine industry but across other livestock systems; more recently studies have attempted to relate different feed categories: blood products from livestock animals (animal by-products), cereal grains (i.e. soybeans, corn, wheat), oil (canola, corn, soybean), forage, pellets (complete compound feed) and straw (bedding material) with the propagation of ASFV (Gordon et al., 2019; EFSA Panel on Animal Health and Welfare (AHAW) et al., 2021; Niederwerder, 2021). Despite the concerns about feed as a route of PRRSV transmission, many uncertainties remain, including the minimal infection dosage required to cause disease, the effectiveness of feed processing such as pelleting, extruding, and roasting and the use of feed additives (Dee et al., 2020; Niederwerder, 2021). In addition, it is important to mention that feed contamination may also occur within the feed mill facility either by contaminated environments, personnel, equipment, birds, or rodents, or even contaminated trailers coming in and out feed mill facilities (Dee et al., 2020; Gebhardt et al., 2021; Niederwerder, 2021), our model does not explicitly consider such uncertainties nor attempt to account for such complexity. In this study we assumed that all feed meals with any amount of animal by-products were still able to cause infection once delivered. In addition, we also assumed that the pelleting process did not eliminate PRRSV contamination, and feed was delivered with enough viral load to cause infection. Even though our results showed a relatively small role of animal by-products (less than 2.6%), there are limitations to our approach and also limited information about the risk and pathways of collateral contamination during or after feed manufacturing. For example, remain unknown the effect of high temperatures and pressure used during pelleting in inactivating PRRSV (Benfield et al., 1992; Van Alstine et al., 1993; Bloemraad et al., 1994; Cochrane et al., 2017), once such data becomes available it will be possible to better assess the role of animal by-products in PRRSV transmission.

### Limitations and final remarks

We identify a number of limitations related to the simulated scenarios and data availability. First, our calculation approaches for the contribution of each transmission route was based on simulations that best calibrated to the observed PRRSV cases in space and time. However, while the sensitivity of the final simulations to detect PRRSV outbreaks was good, results may change once more data about other routes of transmission (e.g. rendering networks) or other relevant disease control interventions are considered such as vaccination programs or on-farm biosecurity. Additionally, the lack of data on transportation vehicles movements and feed delivery of companies B and C limited our ability to examine possible differences and similarities about the contribution of those routes among companies. Even with such limitations, it is important to mention that companies B and C were included in the model because of the relevance of the local transmission in the company-to-company spread of PRRSV, described in details elsewhere (Jara et al., 2020). The 1.5 km radius used to define when a vehicle visited a farm or cleaning station was a limitation for reconstructing the vehicle networks for sites that were 1.5 km from each other, thus in some cases, the contacts were counted towards two or more instead of a single farm (median number of neighbors into 1.5 km = 1). A future alternative to reduce such events would require geographic information of each farms’ feed bins, for example. Additionally, the on-farm model parameters were oversimplified, since we have based those estimations through historical records of PRRSV outbreaks, in which farms with fewer infections were considered to have better biosecurity levels. Although there are several ways that the current version of PigSpread model can be expanded, the inclusions of specific on-farm biosecurity practices and infrastructure (e.g. present of cleaning and disinfection stations) could not only improve model calibration, but to analyze the role of individualized and combined biosecurity on PRRSV dissemination (Sykes et al., 2021). Another important limitation of our modelling work was the lack of information about the vaccination strategies used by each farm, which could have contributed to the probability of new PRRSV outbreaks (Galvis et al., 2021). Finally, we highlight that additional studies are necessary to understand the effect of EVI and seasons on the local transmission, and to evaluate the performance of the gravity model against other transmission methods, such as density kernels, to fit PRRSV outbreak locations. Despite the limitations, this is the first study modelling simultaneously nine routes involved in PRRSV dissemination dynamics over an entire year of outbreak data. This study is unique because it provides the swine industry and regulatory agencies with robust and essential results about the dynamics of between-farms swine disease transmission and the most relevant routes of transmission, offering an unique opportunity to enhance the control of endemic disease and also prepare for future threats (Herrera-Ibatá et al., 2018; Jurado et al., 2019; Brown et al., 2020).

## Conclusion

We expanded a previously developed stochastic PRRSV simulation model (Galvis et al., 2021) to account and quantify the contribution of nine different routes of between-farm transmission, including for the first time the role of animal by-products delivered via feed meals and multiple transportation vehicle networks. Our results demonstrate that transportation vehicle networks have a greater potential to spread PRRSV when compared with the movement of pigs between-farms. In addition, vehicles transporting feed represented the highest risk for PRRSV propagation in comparison with other vehicle networks, connecting around 85% of farms. The temporal network was well represented by its static view for the networks of vehicles transporting feed, and pigs to farms, with causal fidelity values >89%, thus we infer that studies using a static view for vehicle networks are well supported when temporal data is not accessible. Our model demonstrated that pig movements and local transmission remained the main routes of PRRSV transmission regardless of farm types, but vehicles transporting pigs to farms also explained a significant proportion of the farm infections: sow = 20.9%; nursery = 15%; and finisher = 20.6%. As expected, vehicles transporting pigs to markets were more important for PRRSV introduction into finisher farms (3.8%), while vehicles transporting feed showed the highest transmission contribution to sow farms (12%), while the vehicles transporting crew had limited contribution in the propagation of PRRSV regardless of farm types. Finally, animal fat and meat and bone meal delivered via feed contributed to 2.5% and 0.03% of sow farm infections, respectively. Even though we were able to uncover the contribution of by-products and networks of several transportation vehicles in the dissemination of PRRSV, we highlight the need for experimental or observational studies able to measure the viability of PRRSV within feed formulation and the exterior or transportation vehicles. Ultimately, this study provides a better understanding of the role of several transmission routes for PRRSV dissemination, and can provide bases to the swine industry to evaluate and strengthen the surveillance of transportation vehicles and feed delivery to better contain the propagation of PRRSV.

## Acknowledgements

The primary funding support of this project is from the Fat and Proteins Research Foundation. The Morrison Swine Health Monitoring Project is a Swine Health Information Center funded project. Authors would like to acknowledge participating companies, veterinarians and Scott Dee for his useful discussion about the role of feed in the transmission of swine diseases.

## Authors’ contributions

JAG and GM conceived the study. JAG and GM participated in the design of the study. CC coordinated the disease data collection by the Morrison Swine Health Monitoring Program (MSHMP). JAG and GM conducted data processing, cleaning, designed the model, and simulated scenarios. JAG and GM designed the computational analysis. JAG and GM wrote and edited the manuscript. All authors discussed the results and critically reviewed the manuscript. GM secured the funding.

## Ethical statement

The authors confirm the ethical policies of the journal, as noted on the journal’s author guidelines page. Since this work did not involve animal sampling nor questionnaire data collection by the researchers, there was no need for ethics permits.

## Data Availability Statement

The data that support the findings of this study are not publicly available and are protected by confidential agreements, therefore, are not available.

## Supplementary Material

### Section 1. A descriptive analysis of model parameters used in the PRRSV simulated transmission

**Figure S1.**
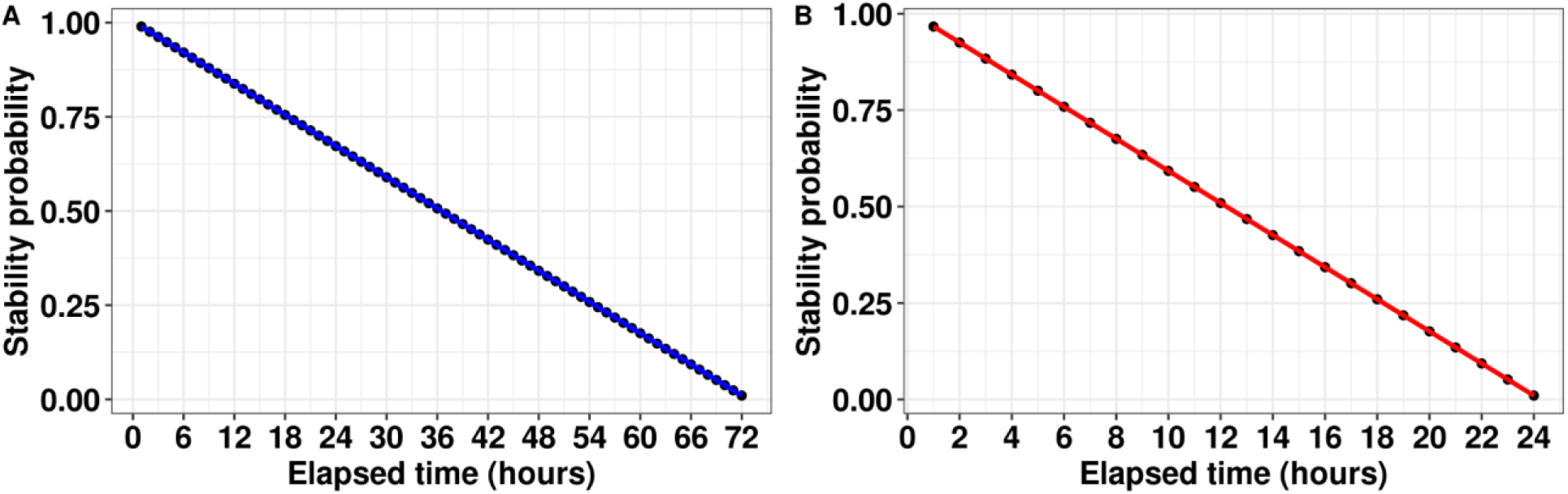
The distribution of PRRSV stability probability on vehicle surface over time. In A), we show the cold season decay which included the months between October to March, and B) warm season decay in PRRSV suitability which included the months between April to September. In summary, here we assumed that PRRSV suitability decreased linearly over time after a vehicle visited a farm, down to zero after 72 hours for cold months and 24 hours for warm months, when PRRSV was no longer viable. This approach was used for all vehicle related contact networks including vehicles transporting feed, pigs-to-farm, pigs-to-market and crew.

**Figure S2.**
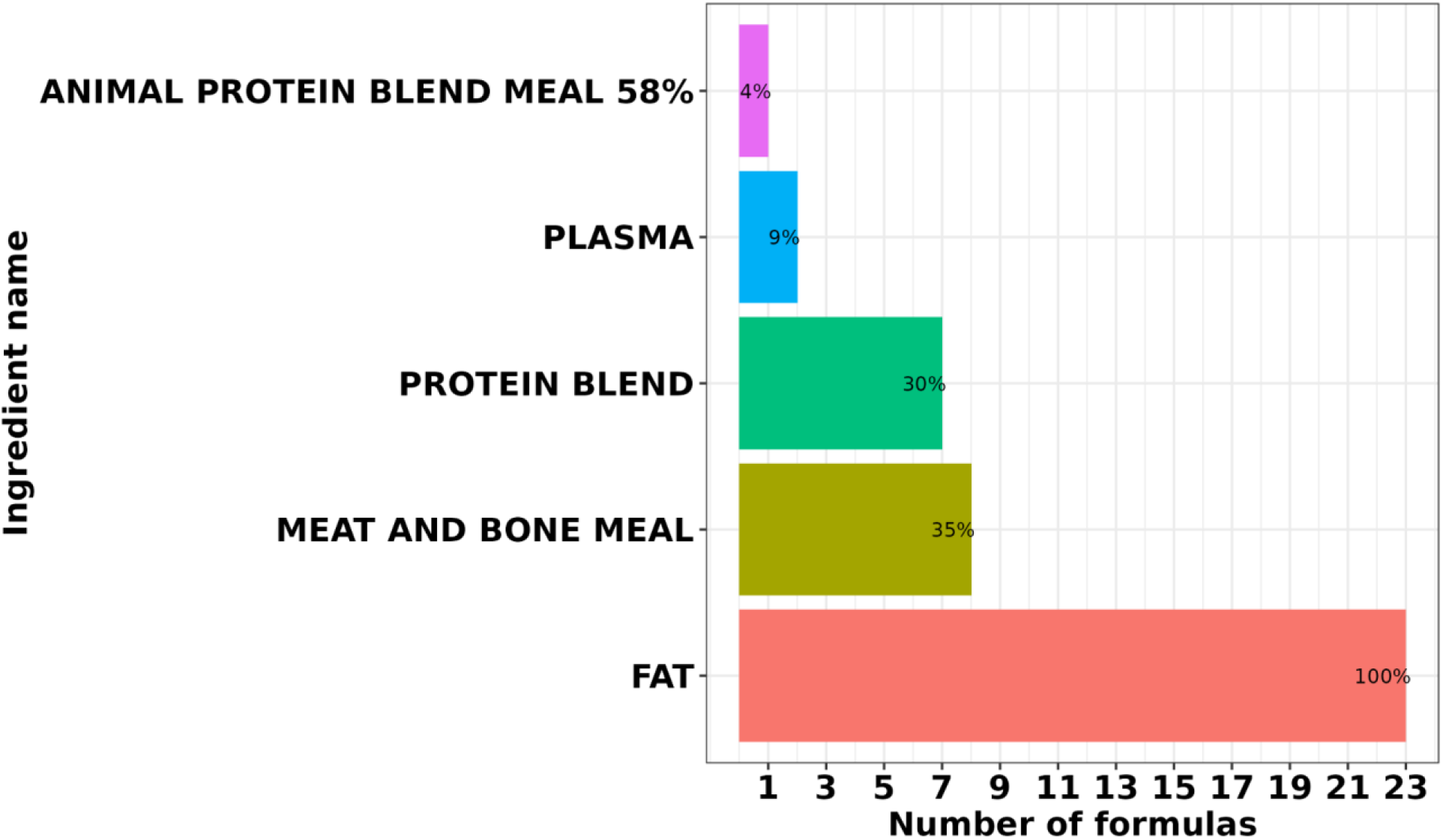
The overall proportion of animal by-products delivered to farms from January 2020 until December 2020. The x-axis shows the number of unique feed formulations and the y-axis each animal by-product utilized by company A.

**Figure S3.**
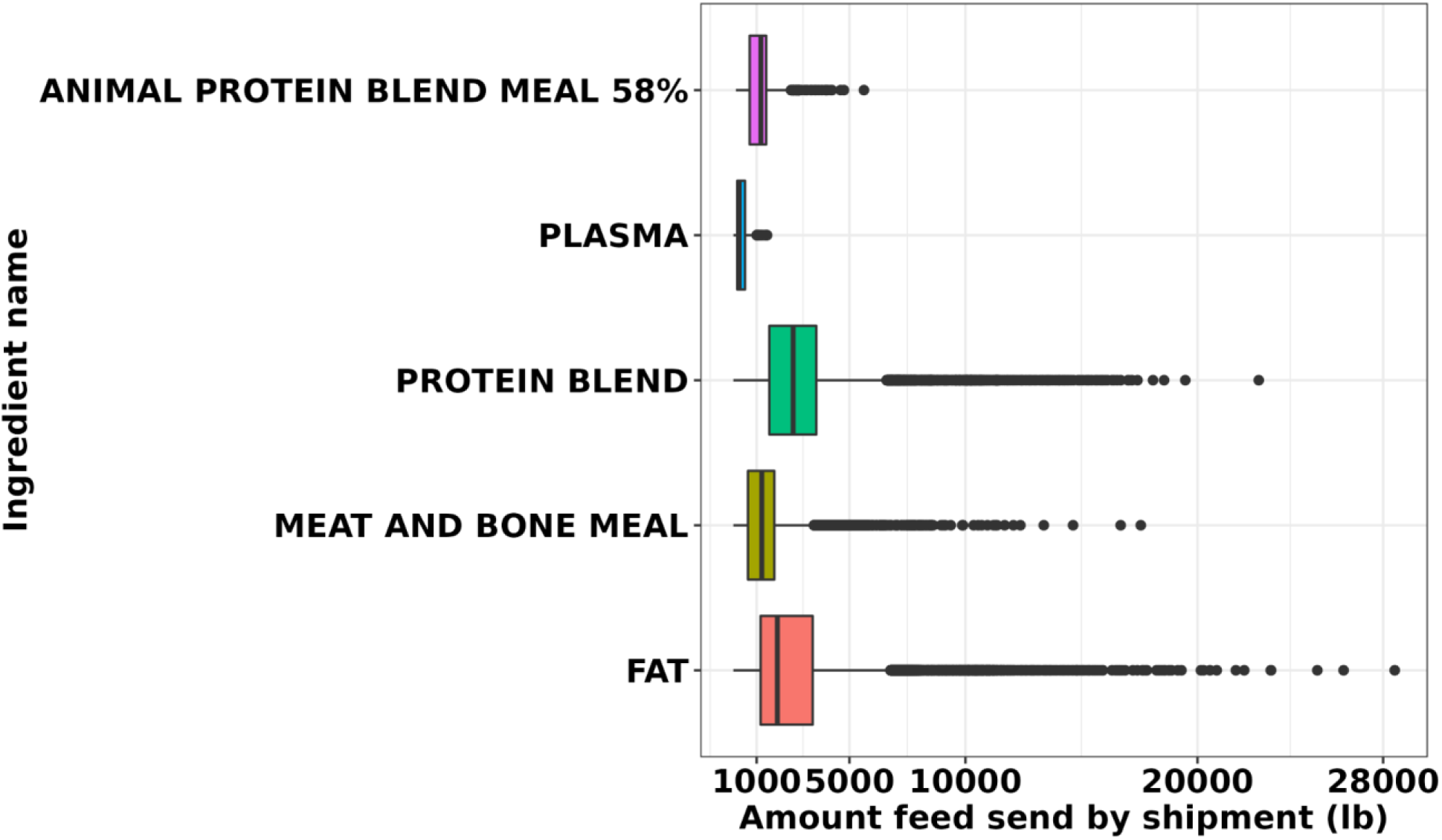
The weight (lb) distribution of animal by-products in shipments of feed formulation delivered to the farms of company A in the study area over from January 2020 until December 2020.

**Table S1.** Terminology and definition from the social network analysis.

| Network terminology | Definition | Reference |
| --- | --- | --- |
| Node | Element of the network representing the farms. | - |
| Edge | Link among two nodes. | - |
| Static network | Once an edge exists between two nodes, it is present for the whole time period. | (Kao et al., 2007) |
| Temporal network | The edges between two nodes only exist at different time steps. | (Lentz et al., 2016) |
| Density | Represent the proportion of edges among nodes in the network that are actually present. | (Wasserman and Faust, 1994) |
| Causal fidelity | Quantify the error of the static representation of a temporal network by comparing the number of paths in the static and temporal networks. | (Lentz et al., 2016) |
| Strongly | It is a subset of nodes for which a *directed* path exists | (Lentz et al., 2016) |
| Connected Component (SCC) | between all pairs of them allowing them to be mutually accessible by following the direction of the links in the network. |  |
| Larger Strongly Connected Component (LSCC) | It is the larger number of nodes in a strong connected component. | (Lentz et al., 2016) |
| In-degree | Number of nodes providing animals to a specific node. | (Wasserman and Faust, 1994) |
| Out-degree | Number of nodes obtaining animals from a specific node. | (Wasserman and Faust, 1994) |
| Betweenness | The frequency at which a node is in the shortest path between pairs of other nodes in the network. | (Freeman, 1978) |
| Ingoing contact chain (ICC) | Subsets of nodes that can reach a specific node by direct contact or indirect contacts through a sequential order of edges through other nodes using the temporal network. | (Nöremark and Widgren, 2014) |
| Outgoing contact chain (OCC) | Subsets of nodes that can be reached by a specific node by direct contact or indirect contacts through a sequential order of edges through other nodes using the temporal network. | (Nöremark and Widgren, 2014) |

To calculate the barrier index (vegetation level, utilized to modulate the probability of local transmission), we used a linear regression, to express PRRSV infected farms from 2020 as a function of the Enhanced Vegetation Index (EVI) and yearly seasonality (spring, summer, fall and winter). We found that PRRSV frequency decreased as EVI increased, with a stronger association in winter and fall seasons (Figure S4). Here we used the regression coefficients to predict weekly PRRSV incidence, which then were transformed into parameter *a*, which was scaled into values between [0, 1], later utilized to modulate the local transmission.

**Figure S4.**
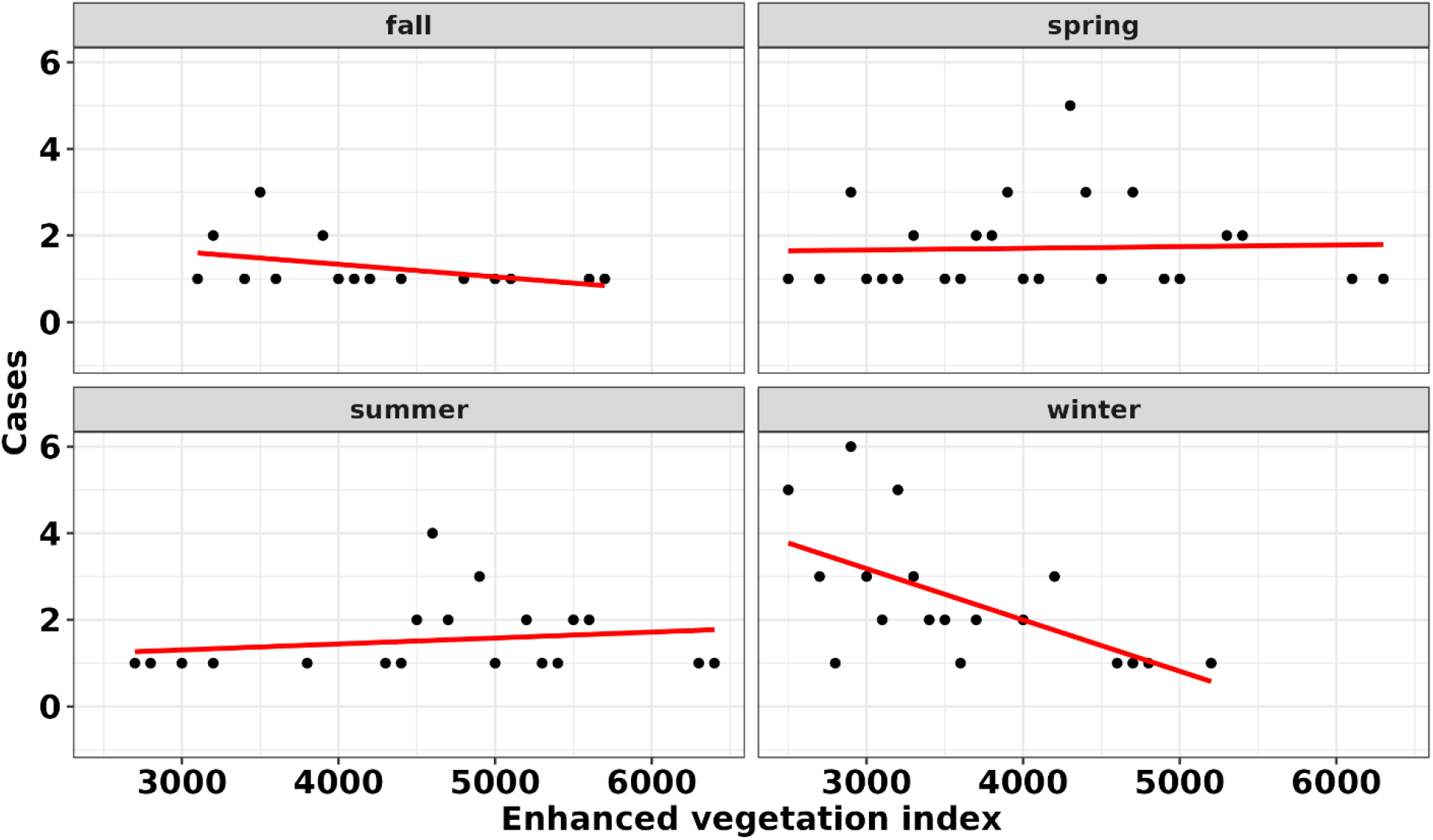
Linear regression of PRRSV infected farms. The y-axis is the number of PRRSV outbreaks and in the x-axis EVI. Fall: coefficient −0.00029, standard error 0.00018, *p*-value 0.12, Spring: coefficient −0.00001, standard error 0.00023, *p*-value 0.96, Summer: coefficient 0.00013, standard error 0.00018, *p*-value 0.46. Winter: coefficient −0.00103, standard error 0.0004, *p*-value 0.02.

**Figure S5.**
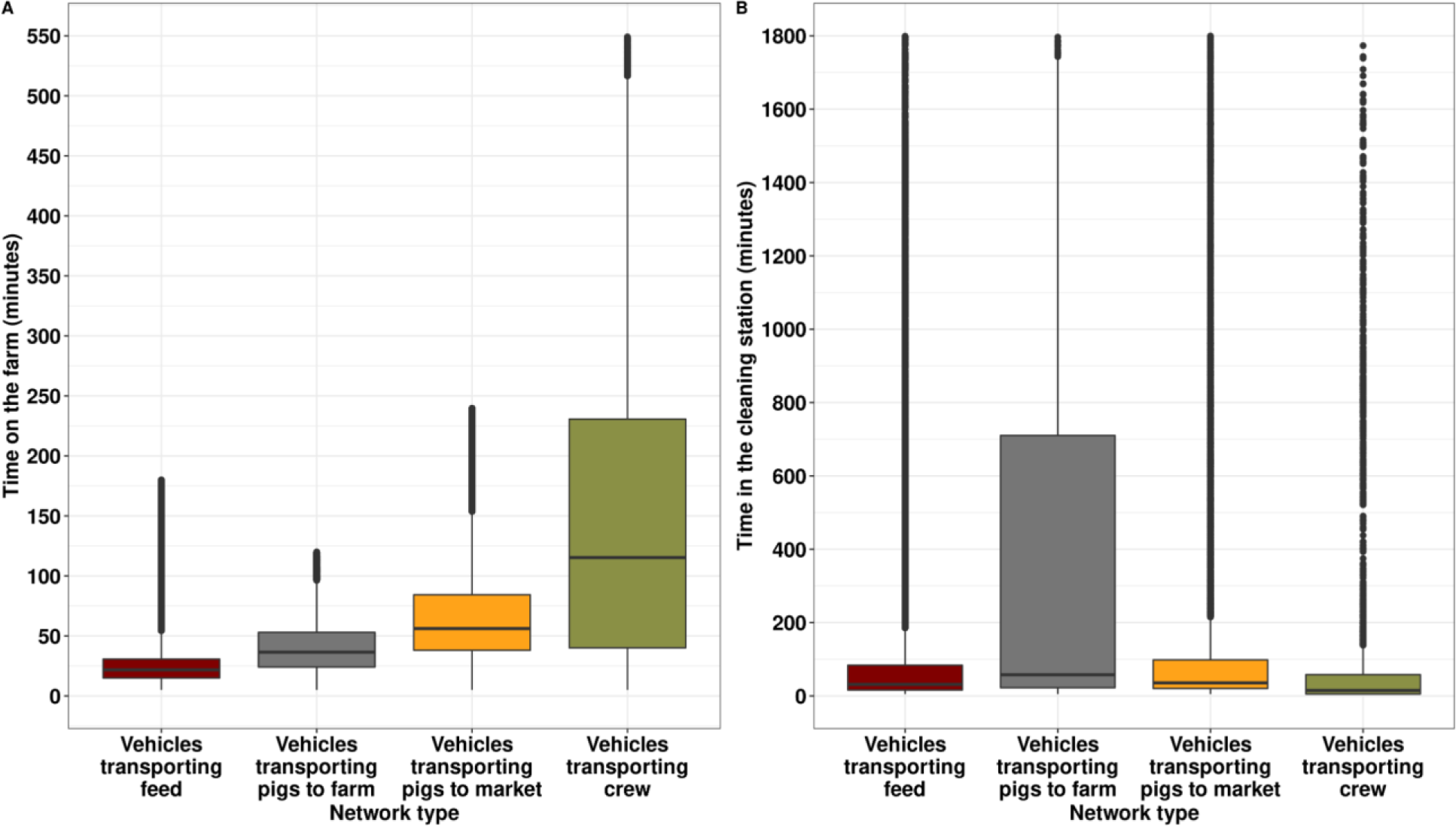
The time vehicles spent on each farm visit. The boxplot shows the distribution in minutes that each vehicle remained within farms premises in A) and at cleaning stations in B).

**Figure S6.**
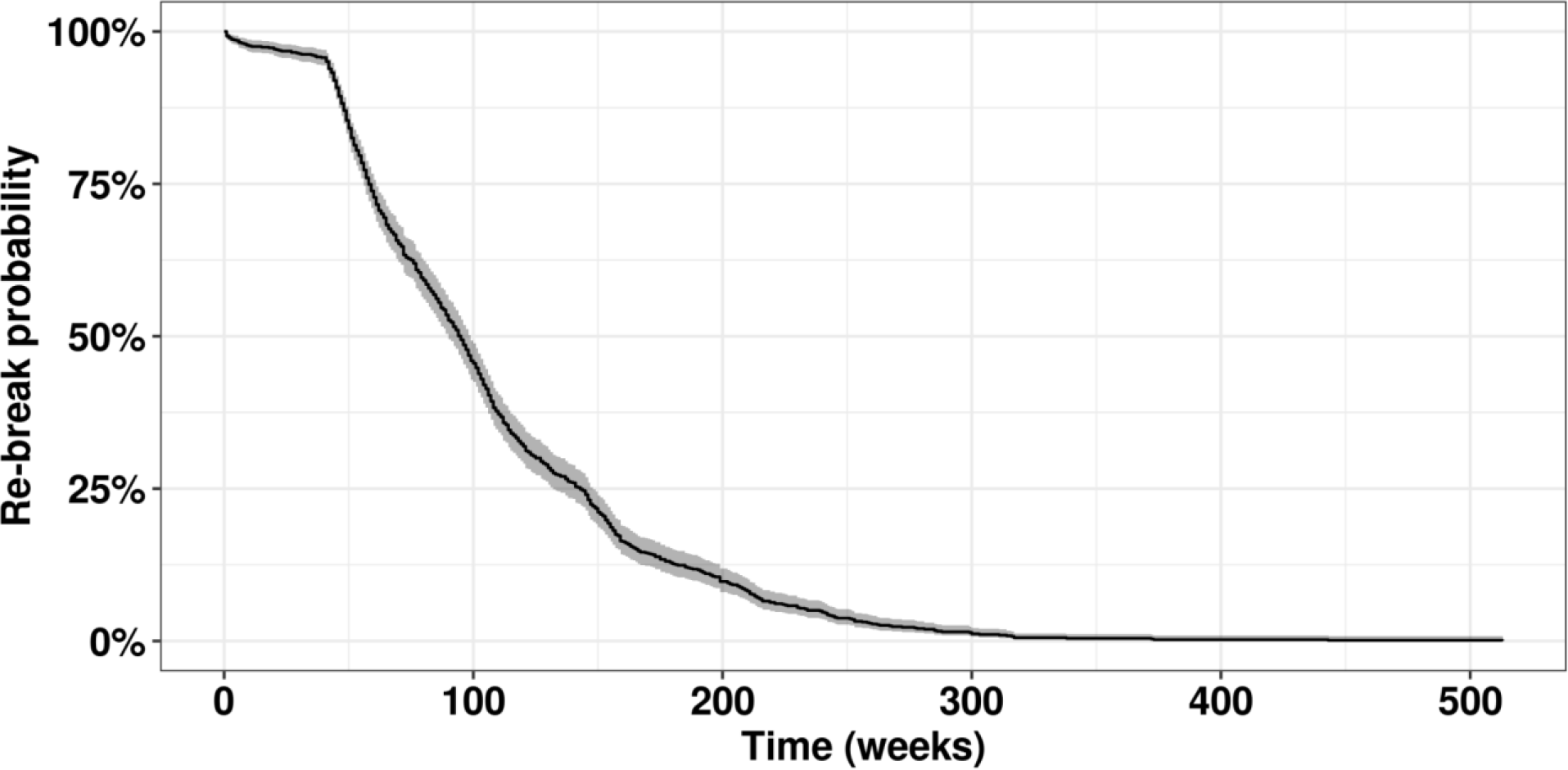
A survival analysis of infected and recovered farms from 2009 to 2019. In this example we show the distribution used for each farm to calibrate the re-break probability

**Figure S7.**
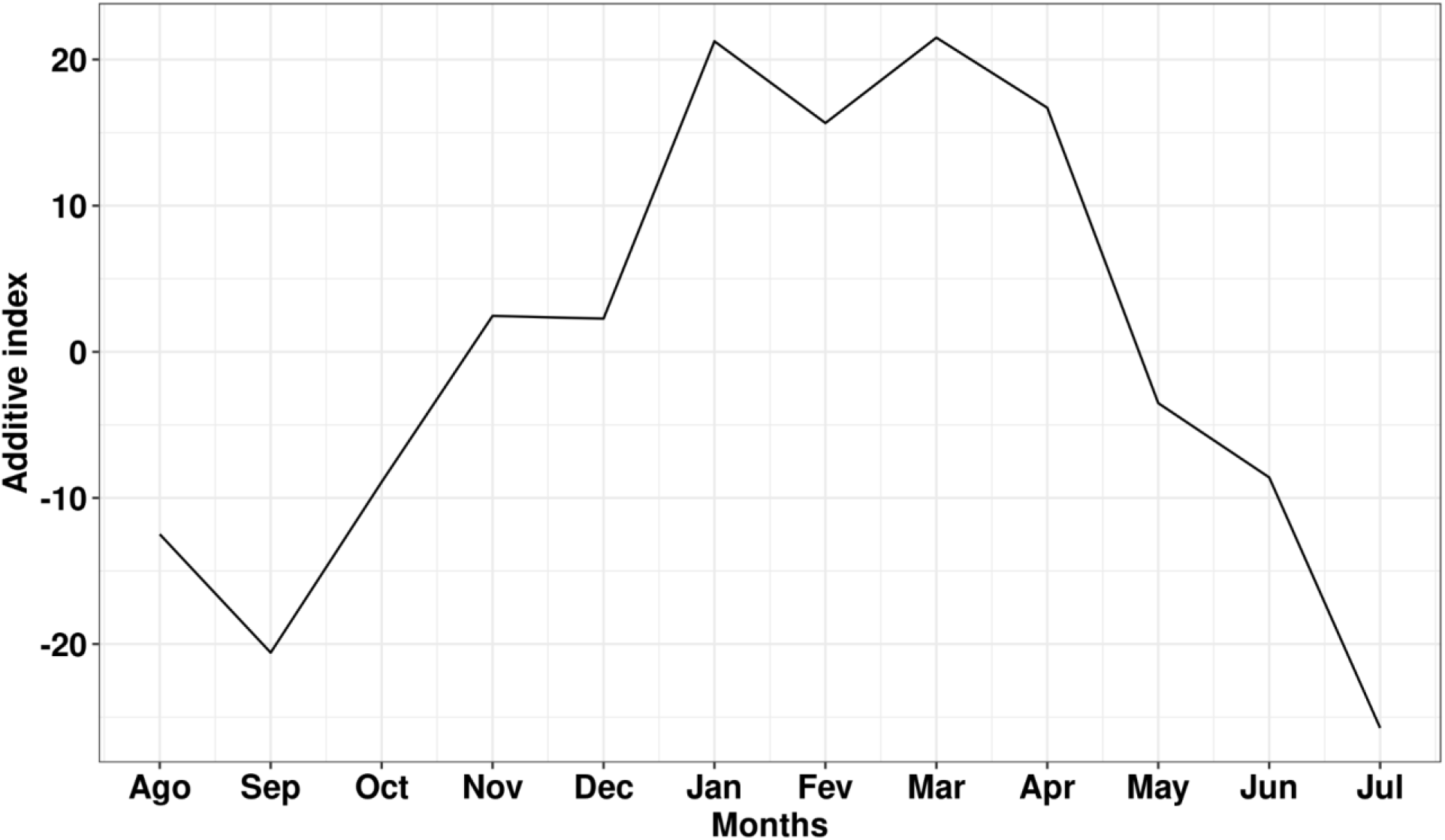
Monthly seasonality index calculated from the frequency of PRRSV, calculated by an additive moving average decomposition derived from analysis of the PRRSV records from 2015 to 2019.

### Section 2: Model calibration and main model outputs

The model is calibrated in two steps, first fitting the frequency of observed outbreaks on time, and second fitting the location of the observed outbreaks. In Table S2 we show the summary statistics used in step 1 of the Approximate Bayesian Computation (ABC) rejection algorithm, where the tolerance interval represents the square error allowed from the simulation to the observed values. In the model calibration, a particle is defined as a set of transmission parameters (Table S3). For the model calibration in step 1, we defined as an accepted particle the set of transmission parameters, whose simulation results are lower than the tolerance interval for all the variables when compared with the observed values. Thus, the range of error accepted (tolerance interval) defines the number of particles accepted in the step 1 of the model calibration. In addition, low tolerance interval values produce particles with similar results to the observed values, but at the same time, the number of particles accepted decreases. Thus, in this study the tolerance interval values were chosen through initial exploratory analysis, selecting the minimum tolerance interval values to accept an adequate number of particles to be analyzed in the step 2 of the model calibration, in this case for each 100,000 particles tested around 20 were accepted in the step 1.

**Table S2.**
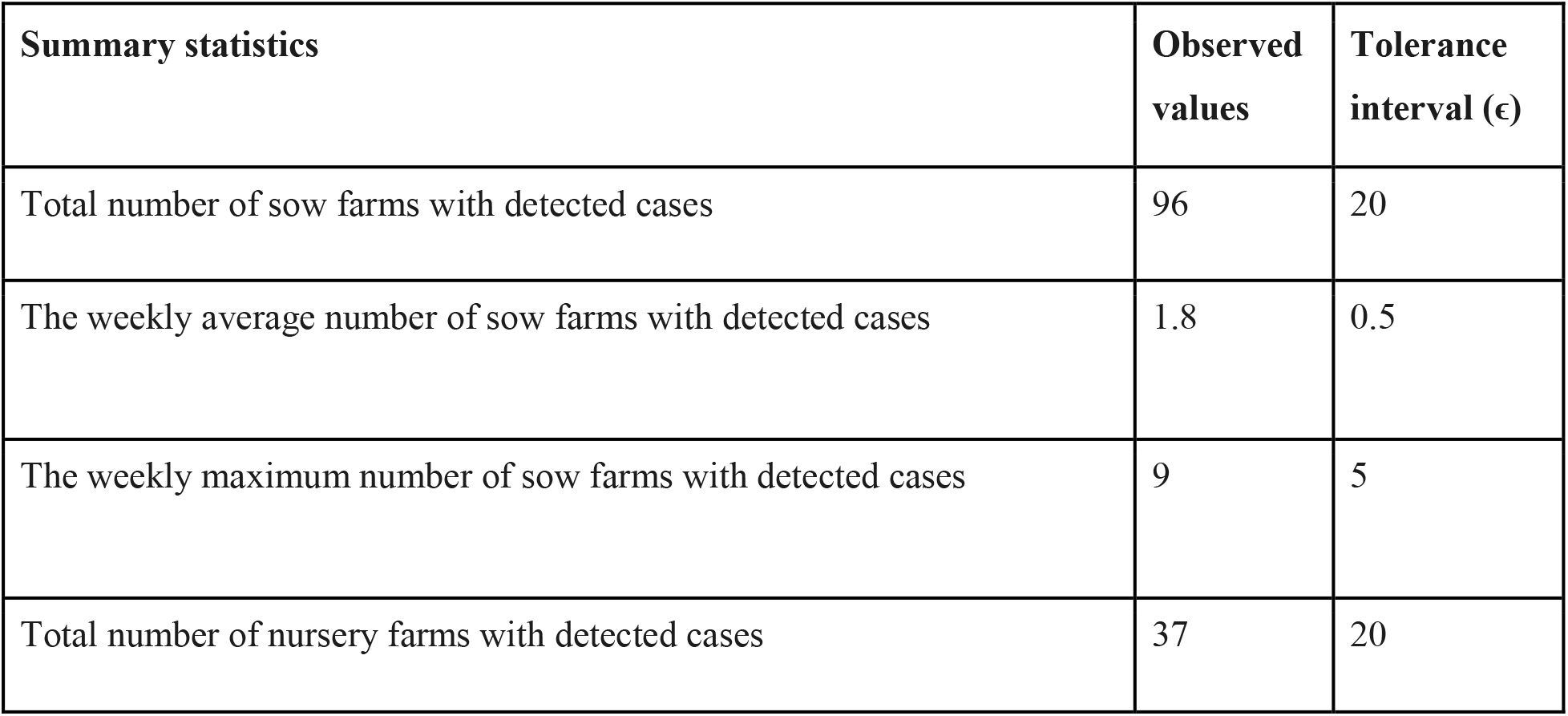
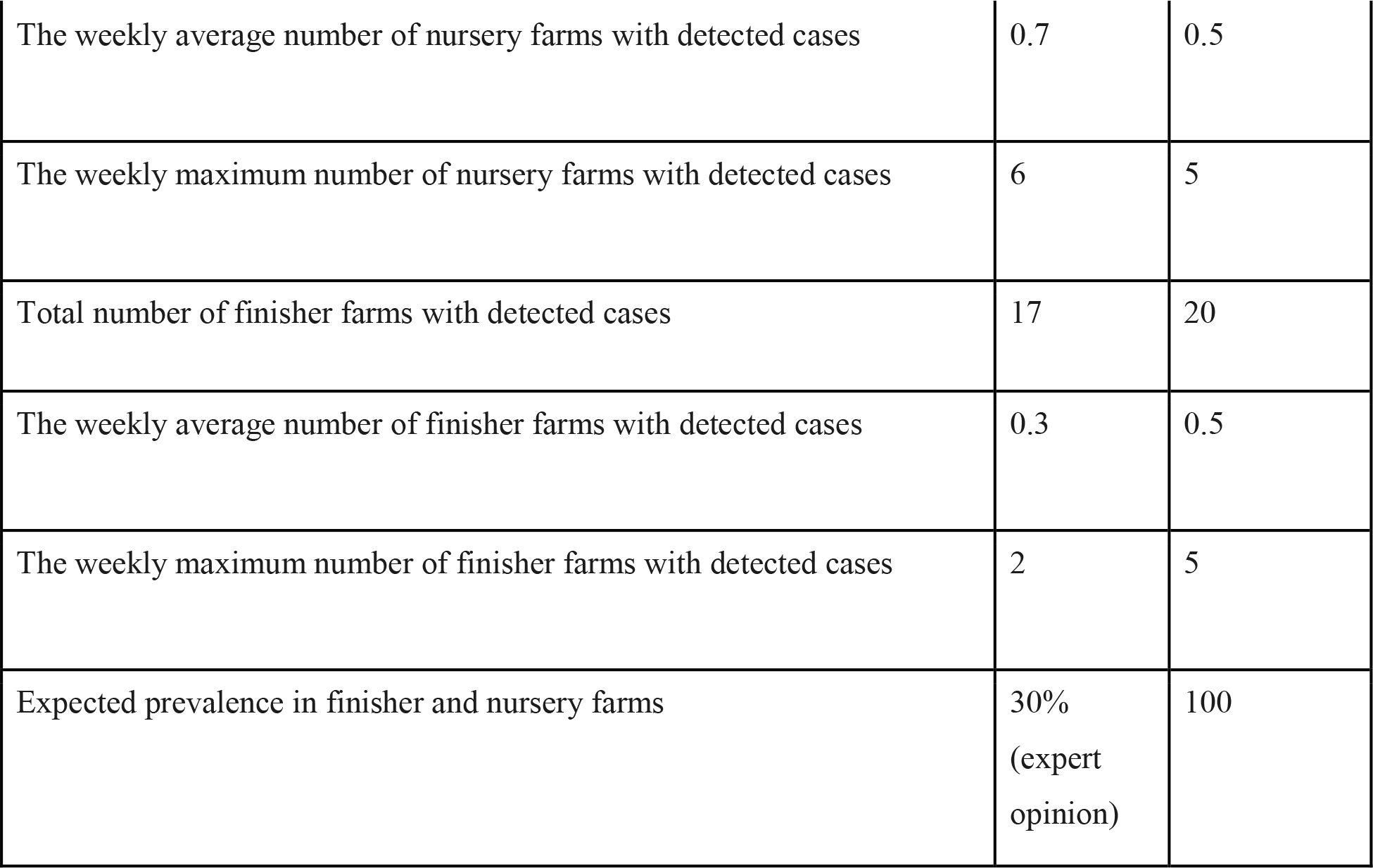
Summary statistics used by the ABC Sequential Monte Carlo rejection algorithm for the model.

To assess the model performance, we evaluated the probability to predict cells (10 x 10 km squares) with true infected cells (cells where at least one sow farm outbreak was recorded) at time *t*. Each sow farm was allocated to a cell in the spatial grid (total of 154 cells in the study area). For each particle accepted in the step 1 model fitting, the risk of each cell was calculated by the sum of times at least one farm within a cell was identified with infected status after 100 simulations; we utilized a percentiles thresholds (r) approach to determine cells at high and low risk, where high risk cells were compared with the true infected cells at time *t*. Subsequently we estimate the model sensitivity and specificity, for all thresholds, as follows:

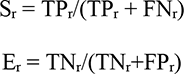

where true positives (TP) was the subset of cells with observed outbreaks and the estimated risk was above the r threshold; false negatives (FN) was the subset of cells with observed outbreaks and the estimated risk was below the r threshold; true negative (TN) was the subset of cells without observed outbreaks and the estimated risk was below the r threshold; and false positives (FP) was the subset of cells without observed outbreaks and the estimated risk above the r threshold.

In step 2 of the ABC rejection algorithm, the sensitivity and specificity were calculated for each particle accepted in the step 1 of model fitting. The particles accepted were those with sensitivity values ≥30% with r = 85th and ≥50% with r = 70^th^ (percentile and sensitivity thresholds values were chosen arbitrarily after authors discussed the minimum performance of the model). The priors for each parameter were drawn from a uniform distribution that ranged between 0 and 1.5 for pig movements transmission rate, 0 and 0.001 for local transmission, the four transporting vehicles and amount of animal fat and meat and bone meal in the feed meals transmission rates, and finally between 0 and 0.01 for re-break transmission rate. These range values were chosen according to model performance to fit the temporal and spatial distribution of PRRSV cases through some test simulations, thus reducing the number of simulations and processing time in the model calibration. It is worth noting that farm’ biosecurity was only included for sow farms, thus there are no parameters representing nursery, finisher or other farms. Finally, here we were not able to describe the number simulation necessary to accept particles, similar how it was described in step 1, because we did not track the number of simulation used in step 2.

**Table S3.** List of transmission parameters used in the simulations and values estimated from the posterior distribution of the 100 particles accepted in the model calibration.

| Model parameter | Symbol | Average values | 95% credible interval* | Details & references |
| --- | --- | --- | --- | --- |
| Transmission rate of pig movements | $\beta_n$ | 0.428 | 0.11-1.29 | ABC fitting |
| Local transmission rate | $\beta_l$ | 0.00055 | 0.0001-0.0009 | ABC fitting |
| Transmission rate of | $\beta_f$ | 0.000014 | 0.0000004-0.00004 | ABC fitting |
| vehicles transporting feed |  |  |  |  |
| Transmission rate of vehicles transporting pigs to farms | $\beta_p$ | 0.00026 | 0.000006-0.0009 | ABC fitting |
| Transmission rate of vehicles transporting pigs to market | $\beta_m$ | 0.00049 | 0.00003-0.0009 | ABC fitting |
| Transmission rate of vehicles transporting crew | $\beta_c$ | 0.00027 | 0.00003-0.0008 | ABC fitting |
| Transmission rate of animal fat in the feed meal | $\beta_a$ | 0.00042 | 0.000002-0.0009 | ABC fitting |
| Transmission rate of meat and bone meal in the feed meal | $\beta_b$ | 0.00042 | 0.000005-0.0009 | ABC fitting |
| Transmission rate of re-break | $\beta_r$ | 0.0045 | 0.0001-0.009 | ABC fitting |
| Farm' biosecurity | H(sow) | 0.57 | 0.02-0.99 | ABC fitting |
| Maximum effective surveillance | L(sow) | 0.95 | - | Expert opinion |
|  | L(nurseries) | 0.048 | 0.016-0.085 | ABC fitting |
|  | L(finisher) | 0.0015 | 0.0001-0.003 | ABC fitting |
|  | L(others) | 0.045 | 0.002-0.086 | ABC fitting |
| PRRSV seasonality | T | Weekly values calculated | - | Figure S7 |
| Average time for PRRSV detection | X0 | 4 weeks | - | Expert opinion |
| Average infectious time sow farms | - | 41 weeks | - | (Sanhueza et al., 2019) |

\*Credible intervals calculated with method equal-tailed interval.

**Figure S8.**
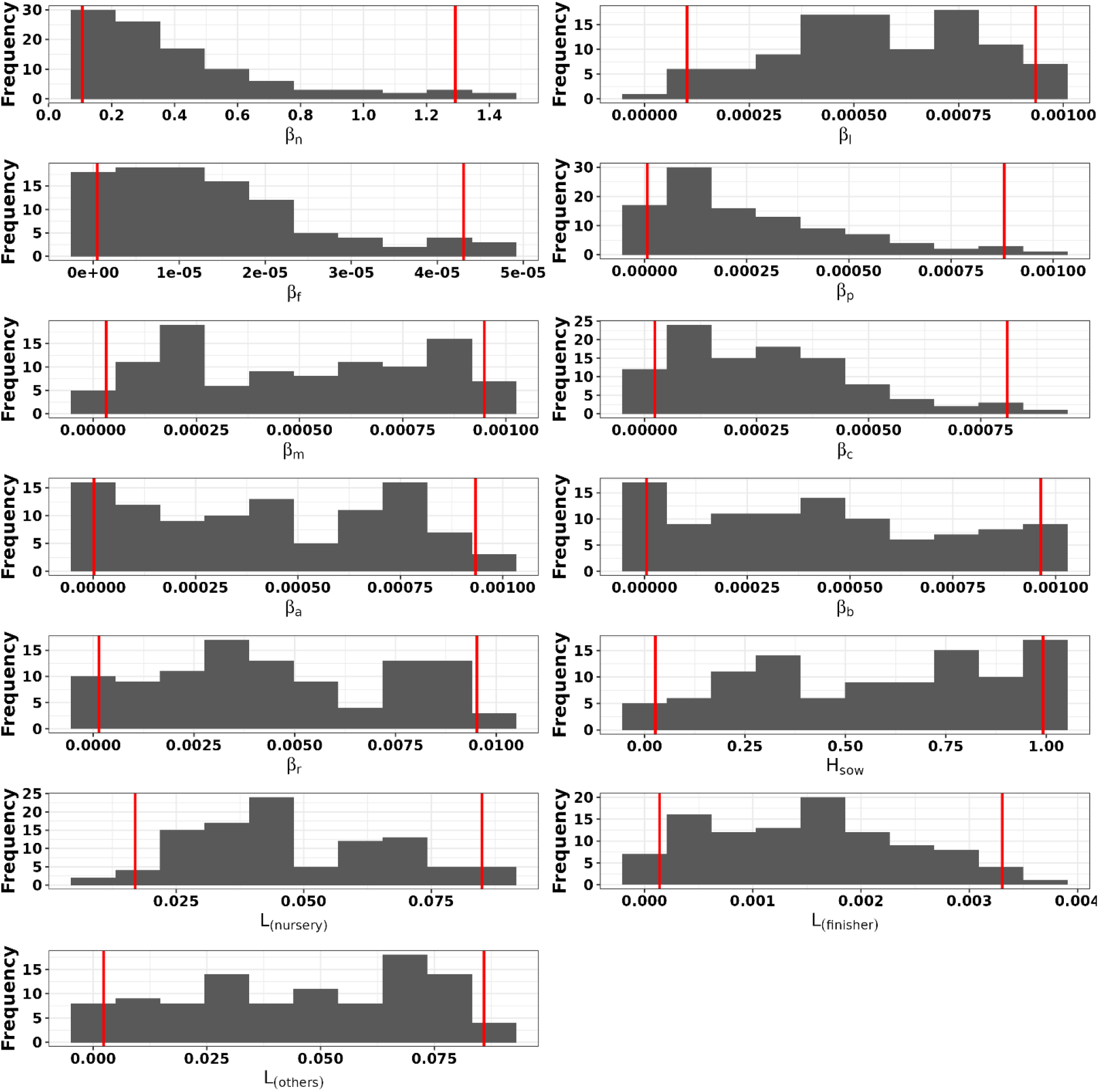
Posterior distribution of the calibrated transmission parameters derived from 100 accepted particles, red lines represent 95% credible intervals calculated through equal-tailed interval method.

**Figure S9.**
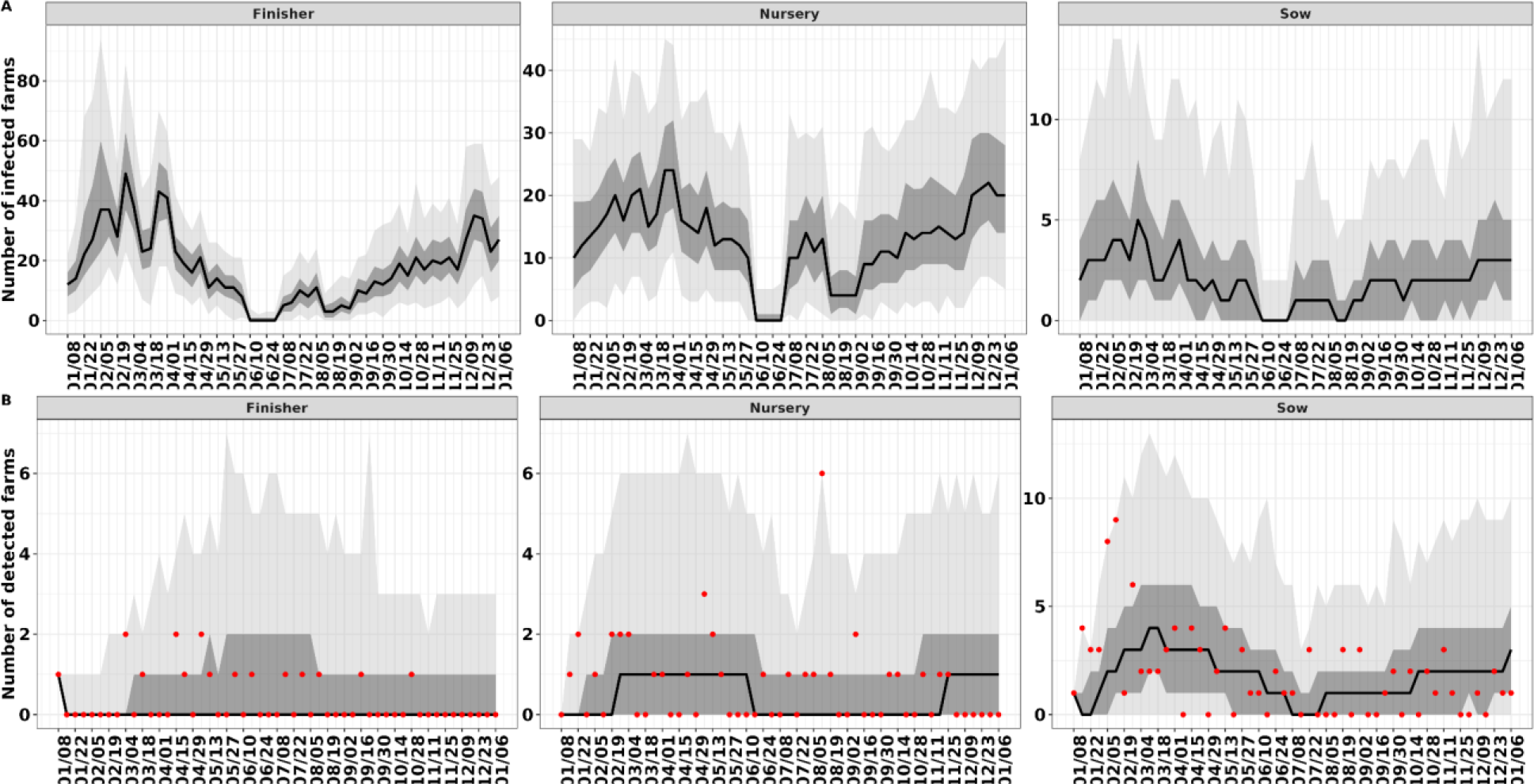
The simulated weekly number of infected farms in A) and infected detected farms (PRRSV outbreaks) in B). The black line represents the median, the dark shade areas represent a 75% credible interval and the light shade areas maximum and minimum generated by the model, and the red dots the frequency of true outbreaks reported in our data. Uncertainty in the estimated model parameters is reflected by 1,000 repeated simulations.

**Figure S10.**
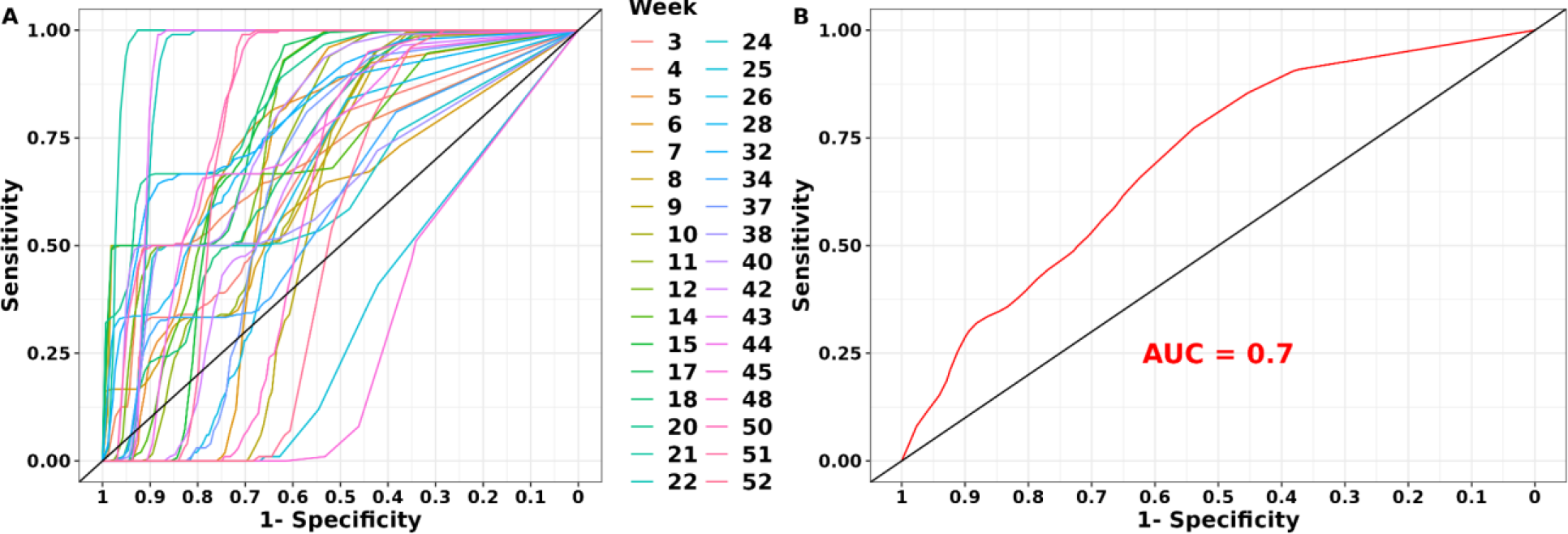
The average sensitivity and specificity for the weekly forecasts in A) and the average of all weeks in B). These values were calculated from 100 model calculations with each model calculation having 100 individual model iterations per predicted week to estimate the spatial location of observed outbreaks

### Section 3: Descriptive analysis of the between-farm pig movements and transportation vehicle movement networks, and the quantity of animal by-product in feed ingredients

**Figure S11.**
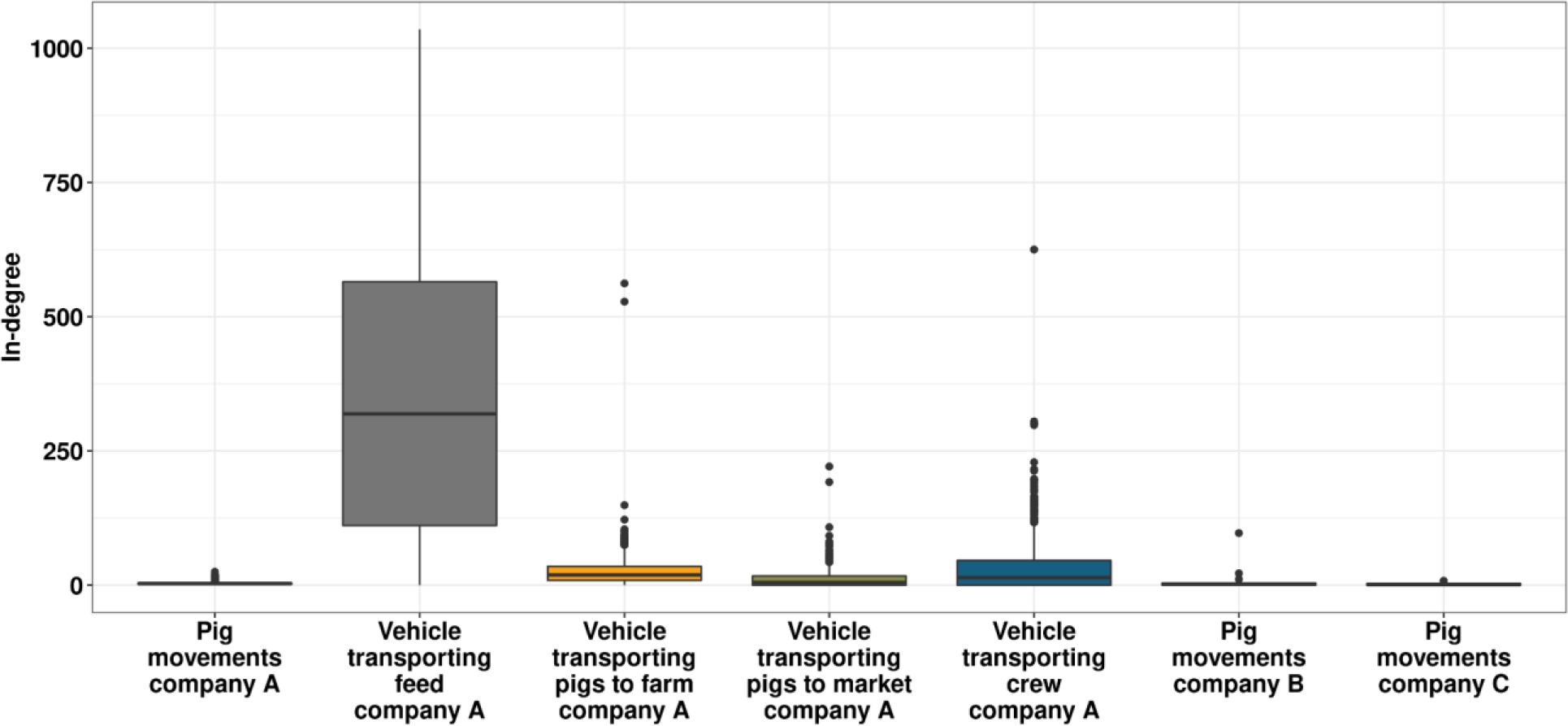
Boxplot with the distribution of in-degree for between-farm pig movements of each transportation vehicle movement networks.

**Figure S12.**
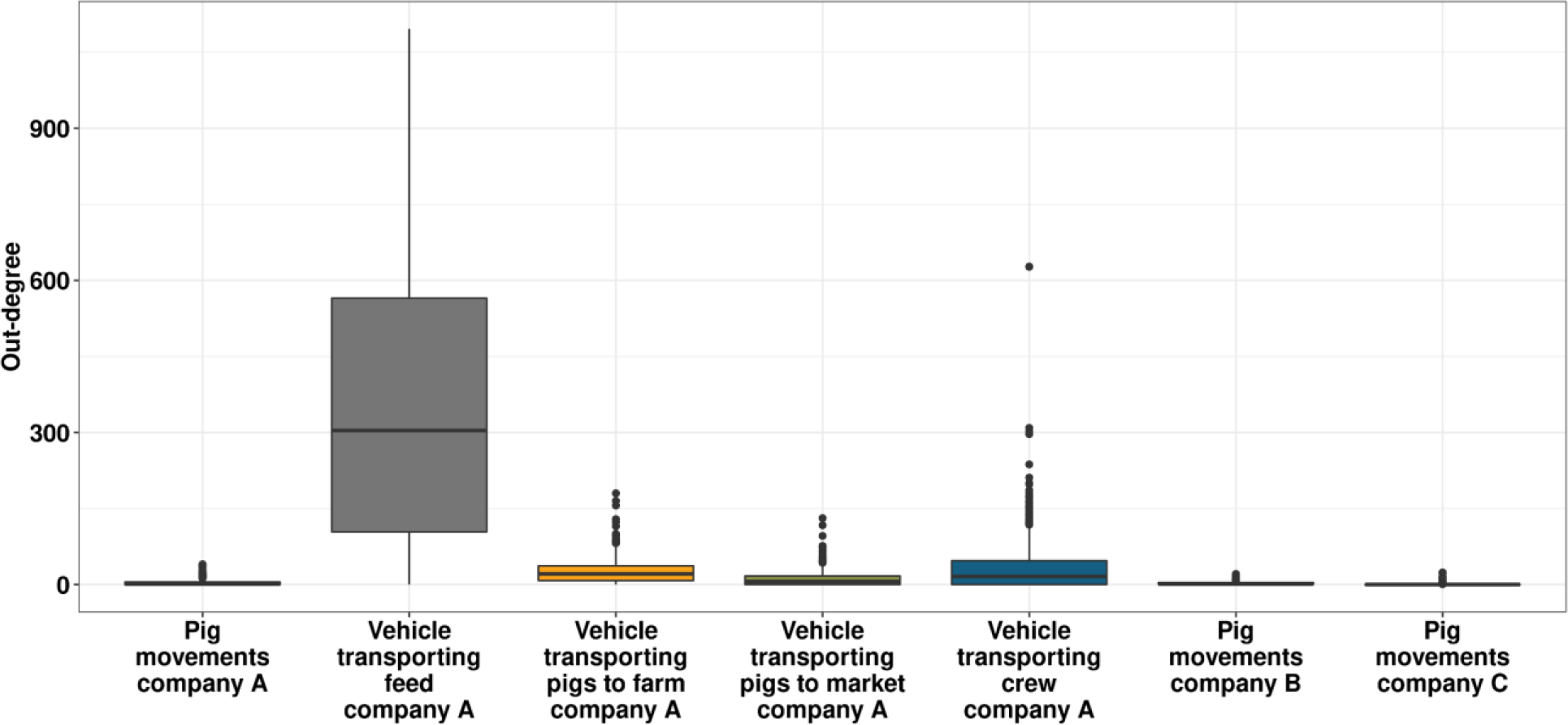
Boxplot with the distribution of out-degree for between-farm pig movements of each transportation vehicle movement networks.

**Figure S13.**
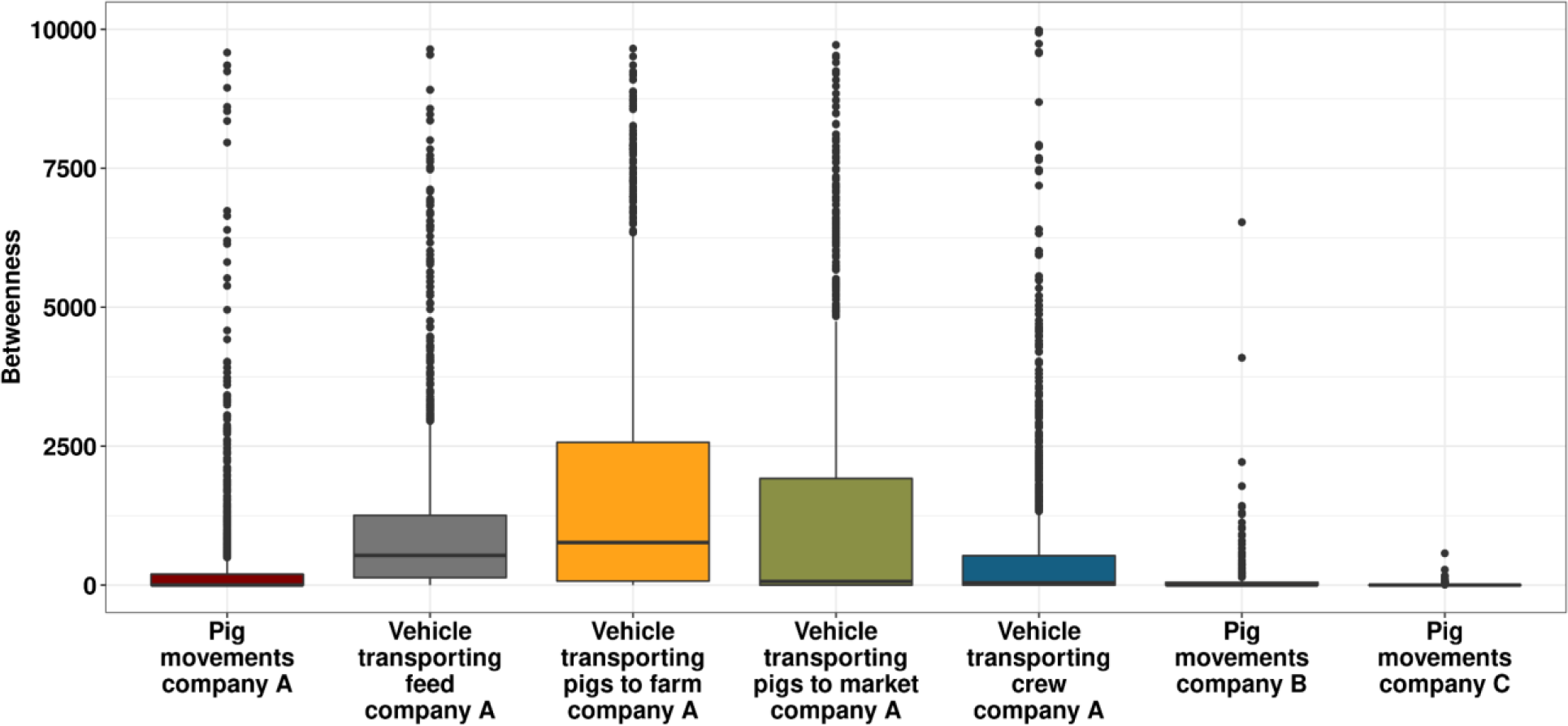
Boxplot with the distribution of betweenness for between-farm pig movements of each transportation vehicle movement networks.

**Figure S14.**
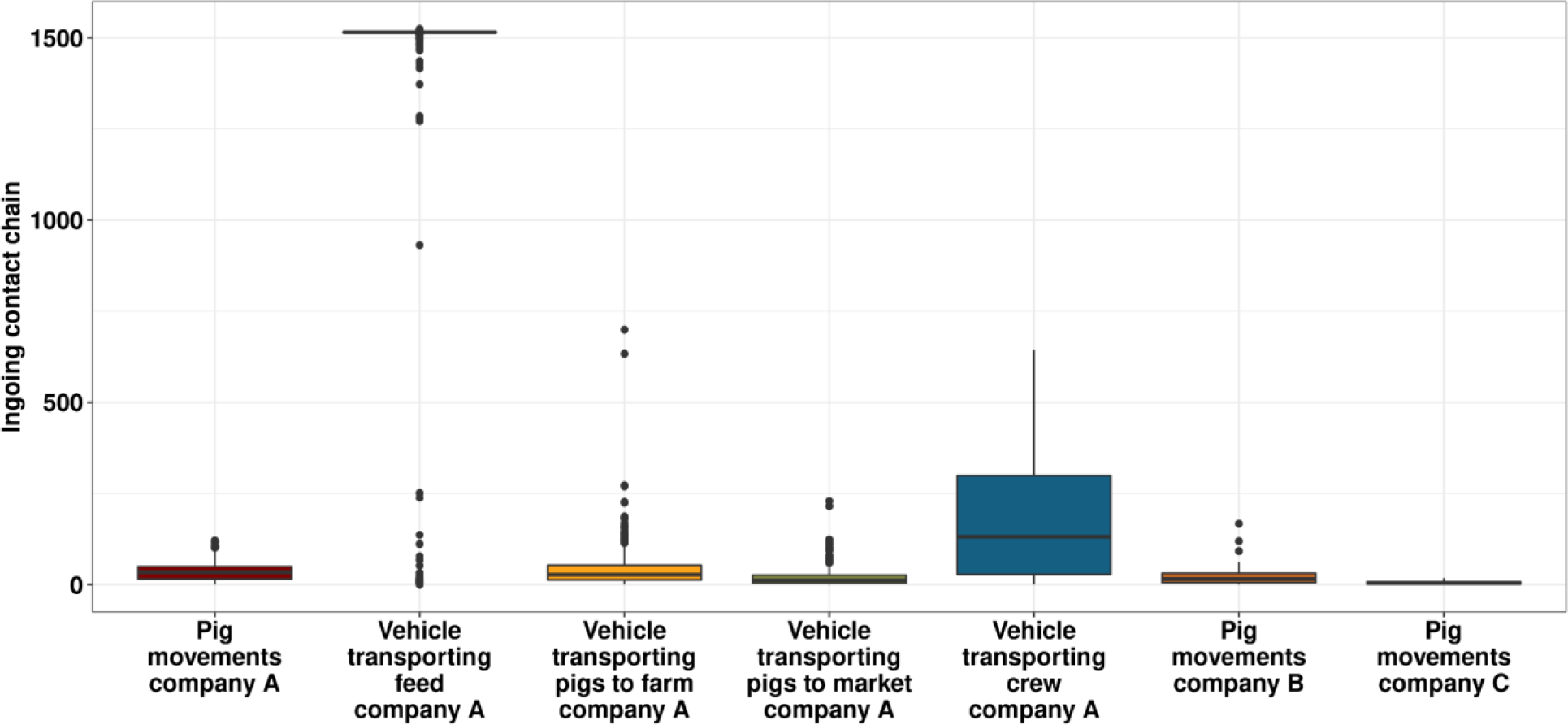
Boxplot with the distribution of ingoing contact chains for between-farm pig movements of each transportation vehicle movement networks.

**Figure S15.**
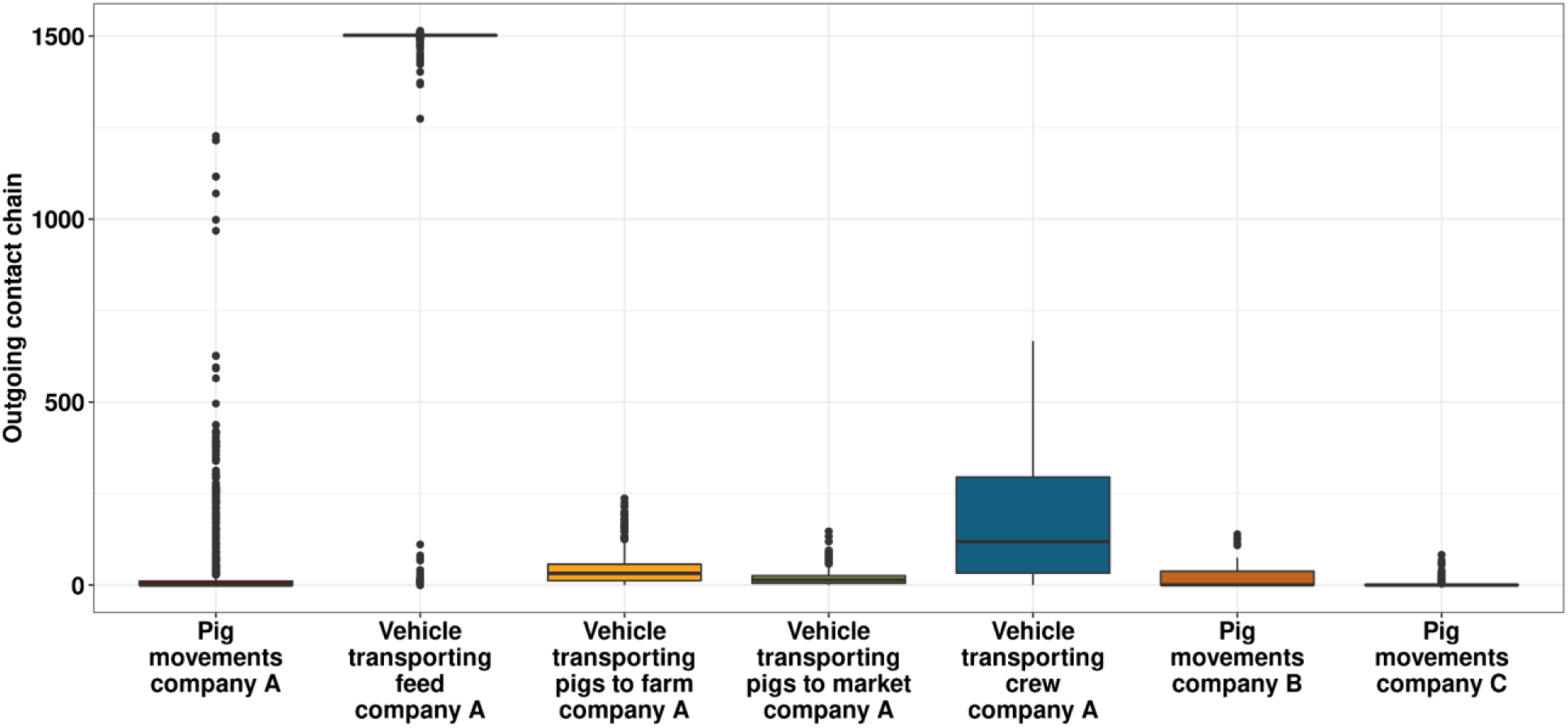
Boxplot with the distribution of outgoing contact chains for between-farm pig movements of each transportation vehicle movement networks.

**Figure S16.**
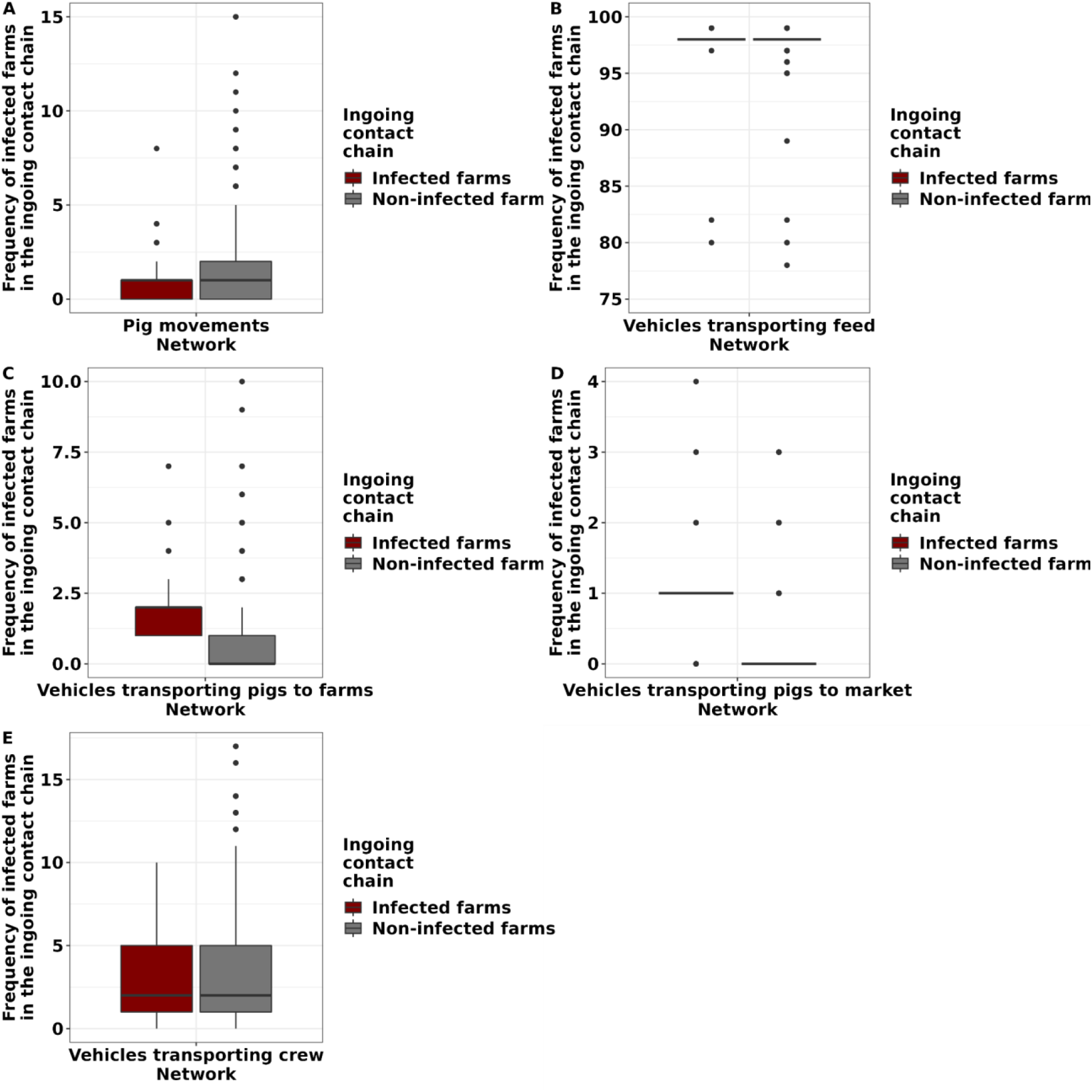
Boxplot comparing the frequency of infected farms in the ingoing contact chain of infected and non-infected farms of each transportation vehicle and pig movement networks. Infected farms are more frequent in the ingoing contact chain of other infected farms for the vehicles transporting feed, pigs to farms and pigs to market (Mann Whitney test *p* < 0.05). It is worth noticing that the statistical differences found among the groups can be influenced by the presence of outliers in the distribution, as well as the large number of farms analyzed (n = 2,294) that increase the test power and make it more likely to reject the null hypothesis.

**Figure S17.**
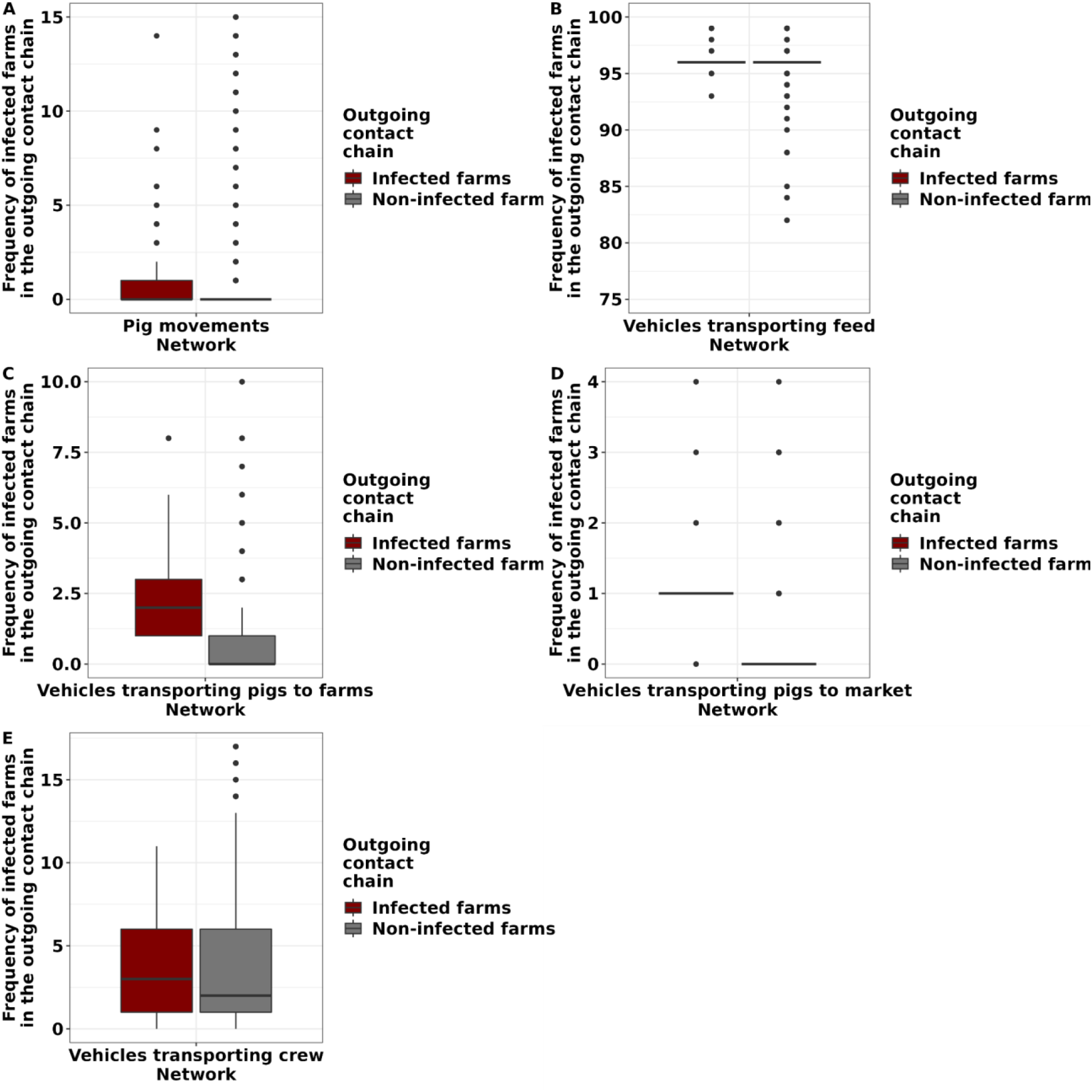
Boxplot comparing the frequency of infected farms in the outgoing contact chain of infected and non-infected farms of each transportation vehicle and pig movement networks. Infected farms are more frequent in the outgoing contact chain of other infected farms for pig movements and the vehicles transporting feed, pigs to farms and pigs to market (Mann Whitney test *p* < 0.05). It is worth noticing that the statistical differences found among the groups can be influenced by the presence of outliers in the distribution, as well as the large number of farms analyzed (n = 2,294) that increase the test power and make it more likely to reject the null hypothesis.

**Figure S18.**
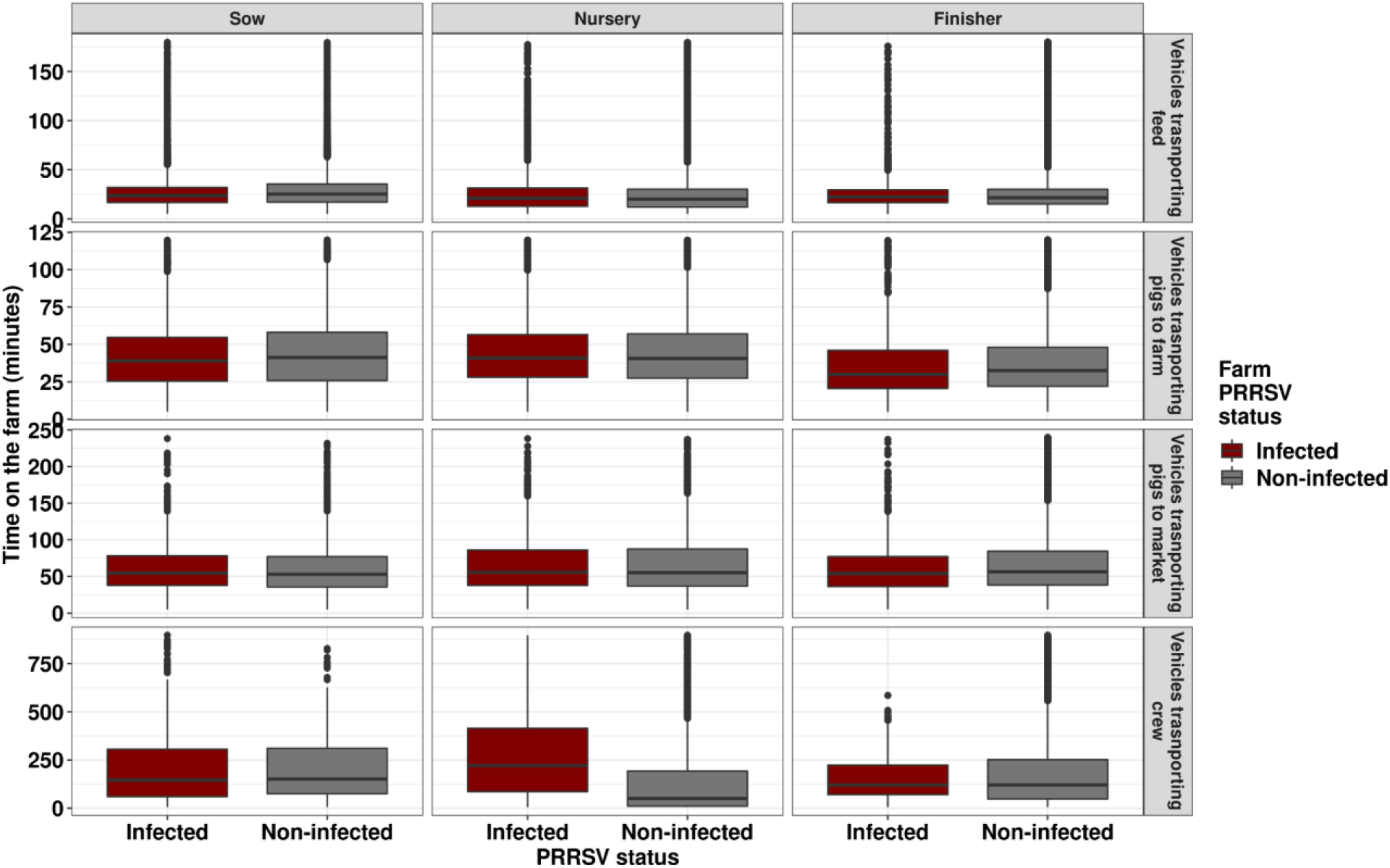
Boxplots compare the time vehicles remain on the farms of infected and non-infected farms for the different transportation vehicles (rows) and production types (columns). Vehicles transporting feed and crew to nursery farms were the only vehicles that showed a higher average of time on infected farms (Mann Whitney test *p* < 0.05).

**Figure S19.**
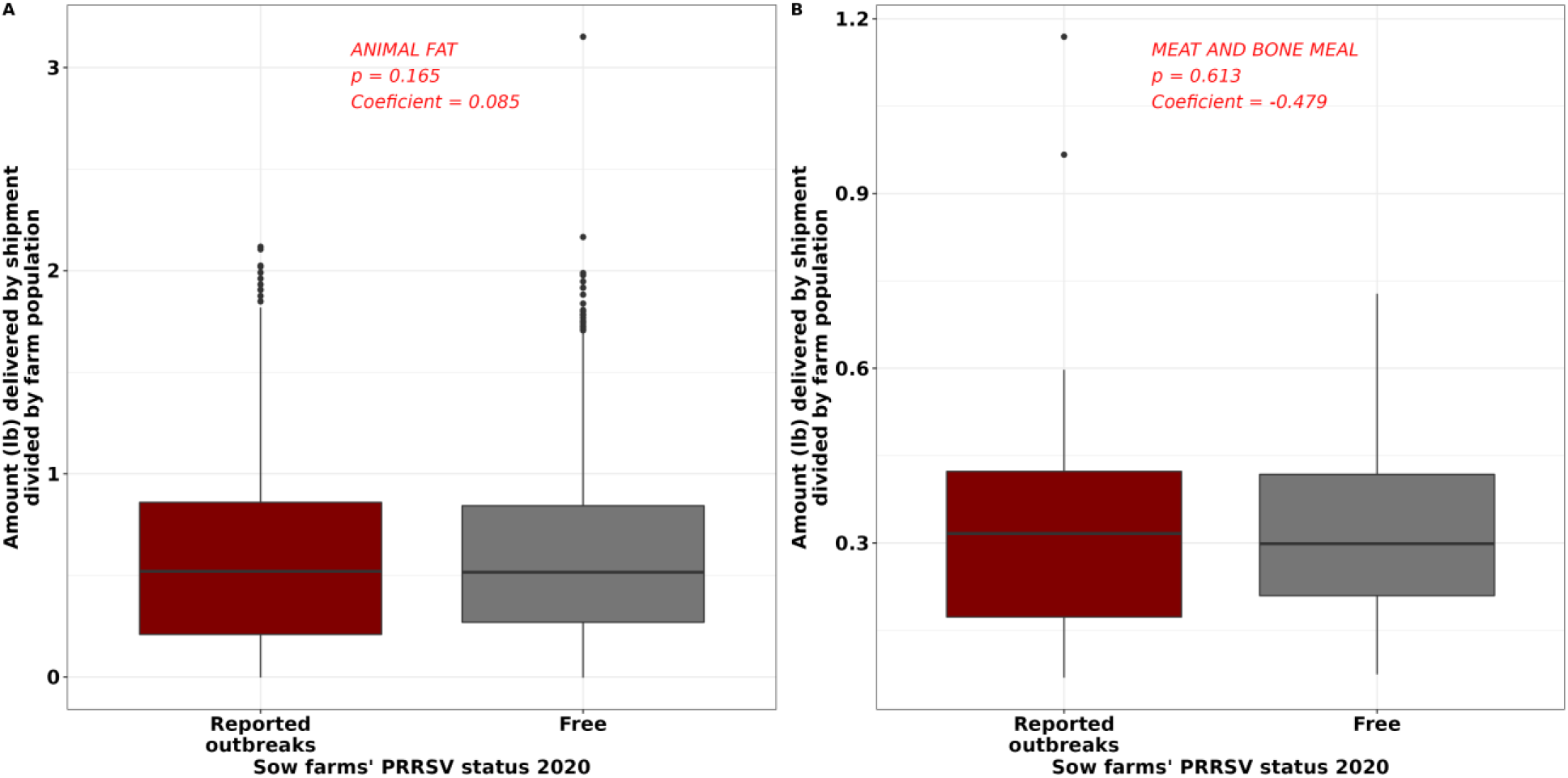
Boxplot comparing the distribution of **A**) animal fat and **B**) meat and bone meal in the feed meal received by the sow farms with and without PRRSV records in 2020 (in red the result from the logistic regression).

**Figure S20.**
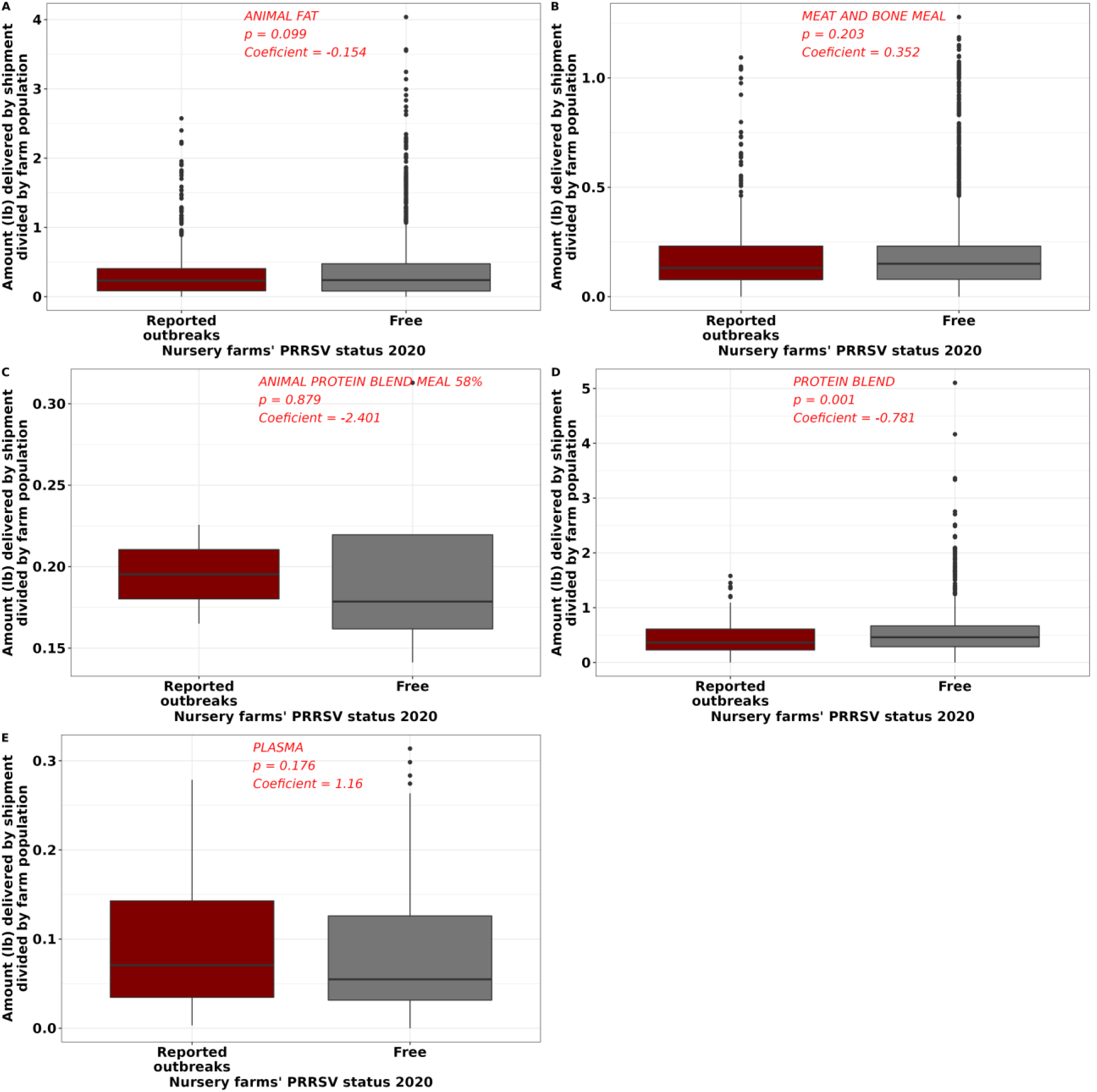
Boxplot comparing the distribution of **A**) animal fat and **B**) meat and bone meal, **C**) Protein blend meal 58%, **D**) Protein blend and **E**) plasma in the feed meal received by the nursery farms with and without PRRSV records in 2020 (in red the result from the logistic regression).

**Figure S21.**
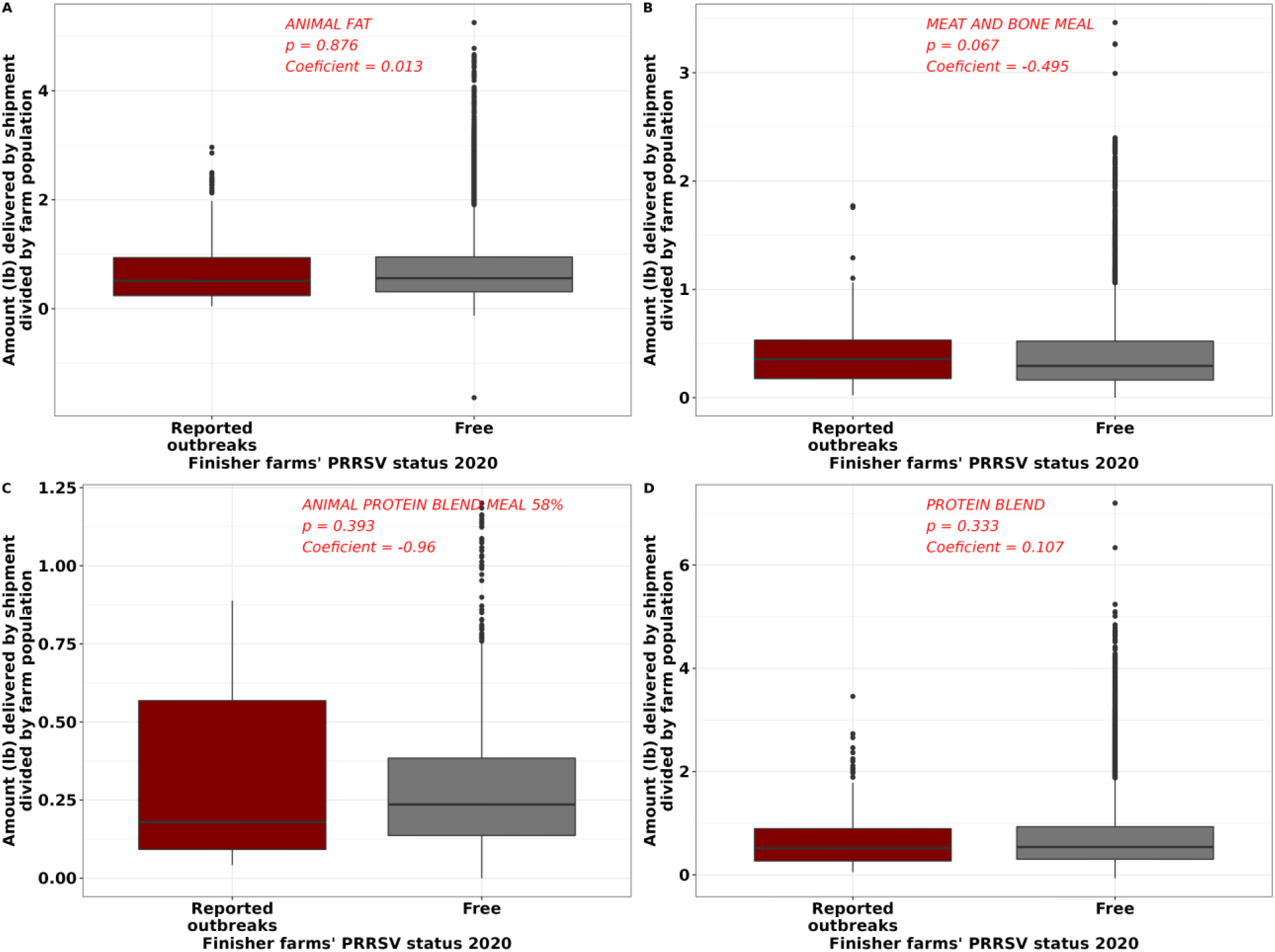
Boxplot comparing the distribution of **A**) animal fat and **B**) meat and bone meal, **C**) Protein blend meal 58% and **D**) protein blend in the feed meal received by the finisher farms with and without PRRSV records in 2020 (in red the result from the logistic regression).

### Section 4:contribution of the transmission routes

For company B, re-break was the main source of farm infections in sow farms with a contribution of 42.6% (95% CI 0%-93%) to PRRSV transmission, followed by local transmission with 28.1% (95% CI 0.8%-88%) and pig movements with 27.3% (95% CI 0%-84%), for nursery 81.6% (95% CI 31%-96%) of PRRSV transmission was related to pig movements and 18.3% (95% CI 4%-68%) to local transmission, while in finishers 51.6% (95% CI 0%-83%) was related to pig movements and 48.4% (95% CI 16%-100%) to local transmission (Figure S22). Finally, for farms of company C, local transmission was the most important transmission route for sow farms contributing with 68.9% (95% CI 8%-97%) of the farm infections, followed by re-break with 26.1% (95% CI 0%-45%) and pig movements 1.2% (95% CI 0%-0%), in nursery 53.2% was related to local transmission (95% CI 8%-100%) and 46.8% (95% CI 0%-92%) to pig movements, and for finishers 78% was related to local transmission (95% CI 31%-100%) and 22% (95% CI 0%-69%) to pig movements (notice that the contribution of transmission routes from the same company and farm type may not reach 100% because for some weeks none of the transmission routes used in the simulations was able to infect farms) (Figure S22).

**Figure S22.**
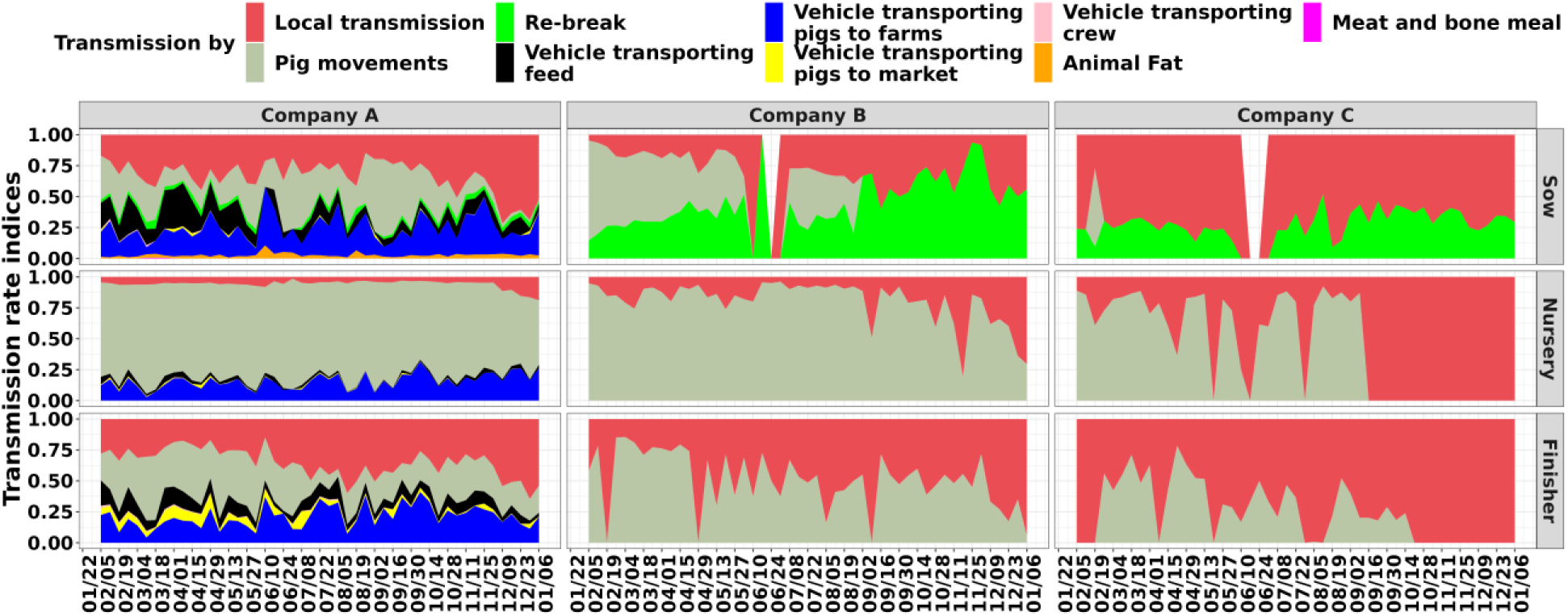
Farm infection contribution for each transmission route of each company (columns) and farm types (rows). The *y*-axis represents the proportion of transmission by each transmission route, while the x-axis shows each week in the simulation. Weekly proportions of transmission were calculated by dividing the number of simulated infected farms for each transmission route by the number of simulated infected farms for the total number of routes combined. White areas represent weeks without farm infections.

